# Highly multiplexed detection of microRNAs, proteins and small molecules using barcoded molecular probes and nanopore sequencing

**DOI:** 10.1101/2022.12.13.520243

**Authors:** Caroline Koch, Benedict Reilly-O’Donnell, Richard Gutierrez, Carla Lucarelli, Fu Siong Ng, Julia Gorelik, Aleksandar P. Ivanov, Joshua B. Edel

**Author notes:** authors contributed equally (Caroline Koch, Benedict Reilly-O’Donnell).

## Abstract

Currently, most blood tests in a clinical setting only investigate a handful of markers. A low-cost, rapid, and highly multiplexed platform for the quantitative detection of blood biomarkers has the potential to advance clinical diagnostics beyond the single biomarker paradigm. In this study, we perform nanopore sequencing of barcoded molecular probes that have been engineered to recognise a panel of biological targets (miRNAs, proteins, and small molecules such as neurotransmitters), allowing for highly multiplexed simultaneous detection. Our workflow is rapid, from sample preparation to results in 1 hour. We also demonstrate that the strategy can be used to detect biomarkers directly from human serum without extraction or amplification. The established method is easily adaptable, as the number and type of targets detected can be greatly expanded depending on the application required.

## Main

The detection of blood serum biomarkers is one of the most common methods for determining diagnosis, prognosis, predicting future disease and monitoring response to treatment. Biochemicals have been used for over 60 years to aid diagnosis of a wide range of conditions, including; cancer^1^, pregnancy^2^ and cardiac disease^3^. Biomarker tests traditionally rely upon the detection of proteins to indicate a condition. However, these tests often lack the specificity to provide clinically useful detail on the pathology. For example, B-type natriuretic peptide (BNP) is used as an indicator for heart failure, but its levels can also be elevated in other related cardiac conditions such as hypertension, cardiac inflammation, and myocardial infarction^4,5^. Combining biomarker assays into a multiplexed platform has clear advantages for the application of multi-biomarker approaches^5^. Such strategies have already been attempted in a number of areas including; Alzheimer’s^6, 7^, amyotrophic lateral sclerosis^8^, cardiovascular disease^9, 10, 11^, chronic obstructive pulmonary disease^12^, infection^13^ and cancers^14, 15, 16, 17, 18, 19^. Many of the methodologies employed rely upon antibody recognition of an epitope and an associated optical readout for each analyte. For example, commercially available systems can detect up to 80 proteins simultaneously^20, 21^. Alternatively, technologies which utilise DNA-based aptamers can detect thousands of proteins in a single sample with a sensitivity down to 125 fM^22, 23^. However, there are limitations to the widespread adoption of such technologies, including large sample volumes, sample labelling and cost.

Clinical diagnostics are now moving beyond protein biomarkers and genetic testing. One example are microRNAs (miRNAs), which are short non-coding RNAs that regulate gene expression^24, 25^. Alterations in miRNA expression have been identified in a wide range of clinical areas: cardiology^26^, hepatology^27^, nephrology^28^, neurology^29^, oncology^30^ and vascular disease^31^. These molecules are detected in the blood primarily through RT-qPCR^32^. However, this method requires multiple steps and relies on signal amplification that may introduce bias in the measurement. Furthermore, the rapid degradation of miRNAs provides a particular problem when considering their use in a clinical setting, as miRNA blood tests are relatively infrequently performed. A rapid miRNA profiling platform would offer the potential to capture short-lived events and perform frequent longitudinal testing.

There is, therefore, a great need to develop new technologies that can perform highly multiplexed detection of various analyte classes, including; nucleic acids, proteins and small molecules. Singlemolecule nanopore sensing offers the ideal platform for performing this task. Nanopores have previously been shown to enable efficient detection of DNA, RNA, proteins, and other molecules^33, 34, 35, 36, 37, 38, 39, 40^, albeit not in a highly multiplexed configuration. Analyte detection by nanopores depends on measuring current fluctuations as charged molecules are electrophoretically driven through a nanoscale aperture. The translocation of a molecule through a nanopore causes a change in the ionic current, which is dependent upon the molecule’s charge, size, and conformation^41^. However, the method generally lacks selectivity. Strategies to address these limitations include chemical modifications of the pore^42, 43, 44, 45^, the use of molecular carriers^46^ and the use of electro-optical methods^47, 48^. These methods are excellent for the detection of molecules at very low concentrations without amplification; however, their throughput and multiplexing ability are limited.

Biological nanopores are advantageous over solid-state devices as they are highly reproducible and can be engineered to manipulate their function^49^. In particular, biological nanopores are useful for DNA/ RNA sequencing, as shown by Oxford Nanopore Technologies (ONT). ONT has commercialised sequencing devices which use biological nanopores arranged in arrays to allow for high throughput, simultaneous DNA/ RNA reads^50^.

This study showcases a new multiplexed analyte detection strategy, combining nanopore sequencing with DNA-barcoded molecular probes. The platform allows accurate demultiplexing of events; simultaneous quantitative detection of at least 40 molecules that can consist of miRNAs, proteins, and small molecules, such as neurotransmitters. The presence of each analyte is determined by the translocation dynamics of each probe as it passes through a nanopore. In pilot experiments, we selected 40 miRNAs and proteins implicated in cardiac disease. The established method is easily adaptable and scalable, meaning the number of detected biomarkers can be extended to cover multiple disease biomarkers at the same time. Due to the simplicity of the experimental protocol and the portability of the platform, we believe that this approach could have an extensive impact on current diagnostics and facilitate the transition to personalised medicine.

### Strategy for the multiplexed detection of analytes

A highly multiplexed detection strategy was achieved by combining nanopore sequencing with barcoded molecular probes that selectively bind to target analytes (Fig. 1a, Supplementary Table 1). Barcoded probes were incubated with target analytes (miRNAs, proteins, small molecules), sequenced with the MinION device (ONT) and subsequently, the presence of target analytes was determined. The barcoded probes consist of three key regions: 1) adapter, 2) barcode and 3) target binding region (Fig. 1B). The adapter is identical for all probes; it includes a motor protein which controls the translocation speed of the barcoded probe to enable sequencing. The adapter region enters the nanopore at its 5’ end, and the 3’ end is capped by a “C3” phosphoramidite spacer molecule. The barcode region is comprised of 35 nucleotides and is capped on the 3’ end by two C3 spacer molecules. The barcode sequence acts as a unique identifier and can have many permutations, with theoretically up to 1.18×10^21^ distinctive arrangements of the nucleotides for the barcode length used in these experiments. The presence of C3 spacer molecules assists in event detection, as these produce higher amplitude current spikes than all other bases. The target binding region can either be a complementary sequence (to bind miRNA or DNA) or an aptamer (to bind proteins and small molecules). The translocation of hybridised probes (target is bound) is slowed since the nanopore geometry does not allow the passage of a double-strand or protein/small molecule to feed through the nanopore (Fig. 1C). The slowed translocation can be observed in the event current signal as a ‘delay’ period. Events can then be sub-classified with or without delay (Fig. 1D), revealing at the single-molecule level the presence of an analyte (Supplementary Fig. 1) whilst the sequenced barcode classifies the analyte being targeted.

**Fig. 1.**
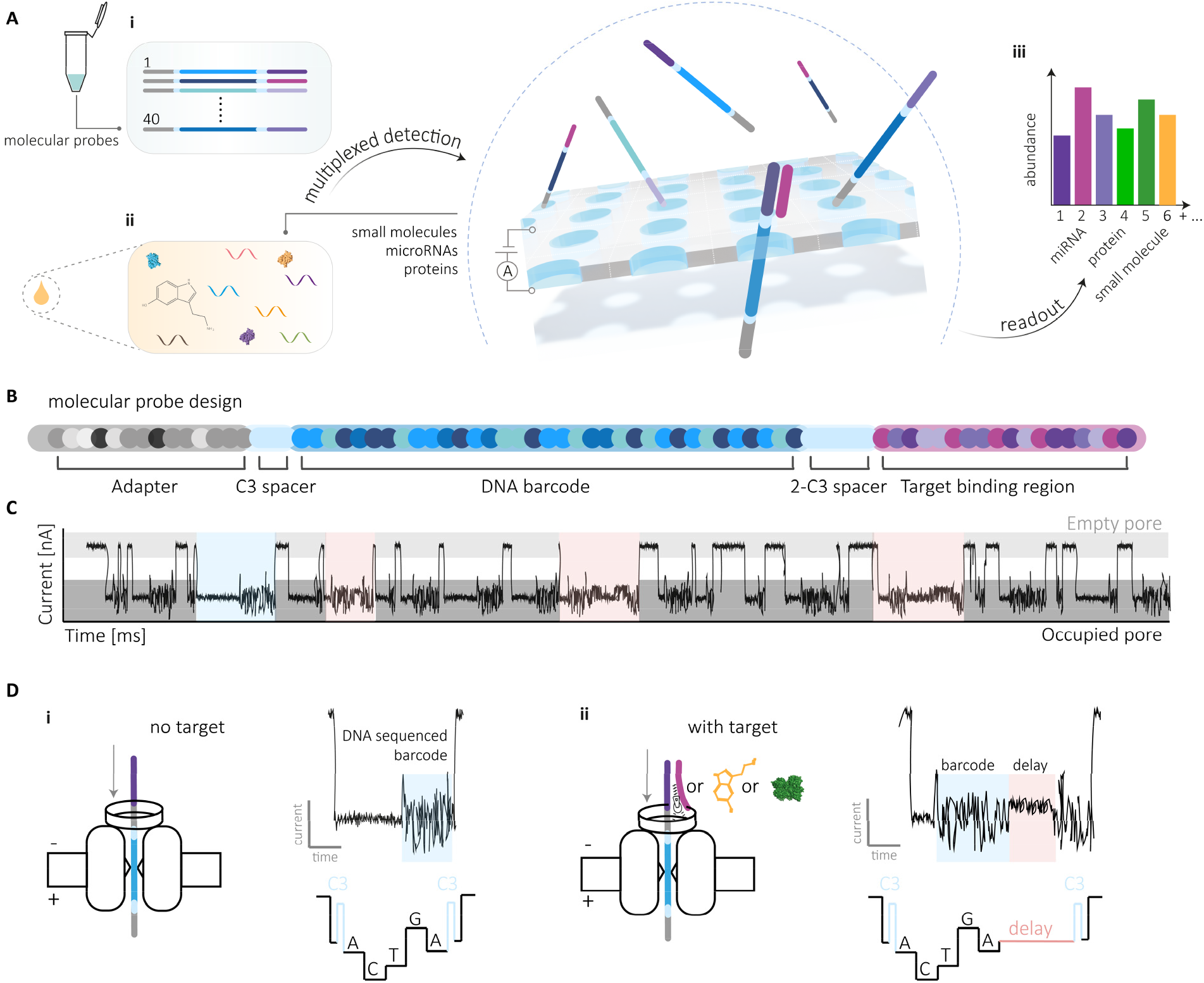
Demultiplexing of 40 miRNAs, proteins and small molecules using barcoded probes for nanopore sequencing. (A) Workflow for detection of miRNAs, proteins, and small molecules. (A, i) Barcoded probes were incubated with synthetic targets or (A, ii) serum samples from healthy participants. Nanopore sequencing was performed to classify the barcode, and (A, iii) the presence of target analytes was determined. (B) Barcoded probe design. The probe consists of a Y-adapter containing the motor protein, a leader strand, and a tether region to facilitate translocation. The barcode region, containing 35 bases, serves as a unique identifier sequence. C3 spacers facilitate the detection of the current signature and downstream processing. The target binding region at the 5’ end of the probe includes either a complementary sequence to a miRNA or an aptamer designed to specifically bind a protein or small molecule. (C) Characteristic current traces. Example events are highlighted for barcode only (blue) and barcode events with bound target analytes (pink). (D, i) Basecalling and analysis of events from barcoded probes result in current traces without delay. (D, ii) Basecalling and analysis of events from barcoded probes with analyte-bound results in current traces with a delay.

To establish the multiplexed platform, we designed 40 unique barcoded probes to detect miRNAs implicated in cardiac disease. The accuracy of barcode basecalling and classification was determined in experiments without target analytes (Fig. 2). Events were sequenced and aligned against a barcode library with known barcode sequences (Supplementary Table 1). For example, the alignment score of a unique barcode, ‘barcode 38’, is shown in Fig. 2a. The normalised alignment score of events for the true sequence was significantly higher (ANOVA, alpha= 0.0001) compared to all other barcode sequences in the library (Fig. 2a, Supplementary Fig. 2a, Supplementary Table 2). However, alignment scoring was not suitable in multiplexed experiments due to the occurrence of false positives. To address this, a series of thresholds were tested including 1) the number of mismatches, 2) *x* mismatches in the first *y* bases, 3) number of aligned bases, and 4) sequence beginning with the bases “GGG” (Fig. 2b). It was found that reducing the total number of mismatched bases worked poorly on its own, as it reduced the number of both true positive and false positive events (AUC= 0.414). We observed that mismatches at the start of the sequenced event were particularly indicative of poorly resolved events. Consequently, a threshold based upon allowing *x* mismatches in the first *y* bases was found to be moderately effective in removing unwanted events (AUC= 0.735). Increasing the total number of aligned bases was the second-best applied threshold (AUC= 0.828). Requiring the sequenced events to start with Gs (as this is common among all barcoded probes) proved to be the most efficient at separating true and false events (AUC= 0.846). Finally, we combined each criterion to determine the optimum threshold: ≤5 mismatched bases total; 1 mismatch in the first 10 bases; ≥15 bases aligned in total; sequence starts with “GGG”. When combining all thresholds, there was no improvement in sensitivity (AUC= 0.805); however, there was a marked reduction in the false positive rate. The introduction of this optimised threshold resulted in an accuracy of >95% in the alignment of basecalled events for single barcoded probe experiments (Fig. 2c). To determine the sensitivity of the platform, we tested a barcode against two further probes with one and two mismatched bases (barcode 8 vs barcode 8 with one mismatch and barcode 8 with two mismatches). In a multiplexed experiment with all three of the barcoded probes present, the alignment score for the correct barcode was significantly higher than the alignment scores of the other sequences (Fig. 2d).

**Fig. 2.**
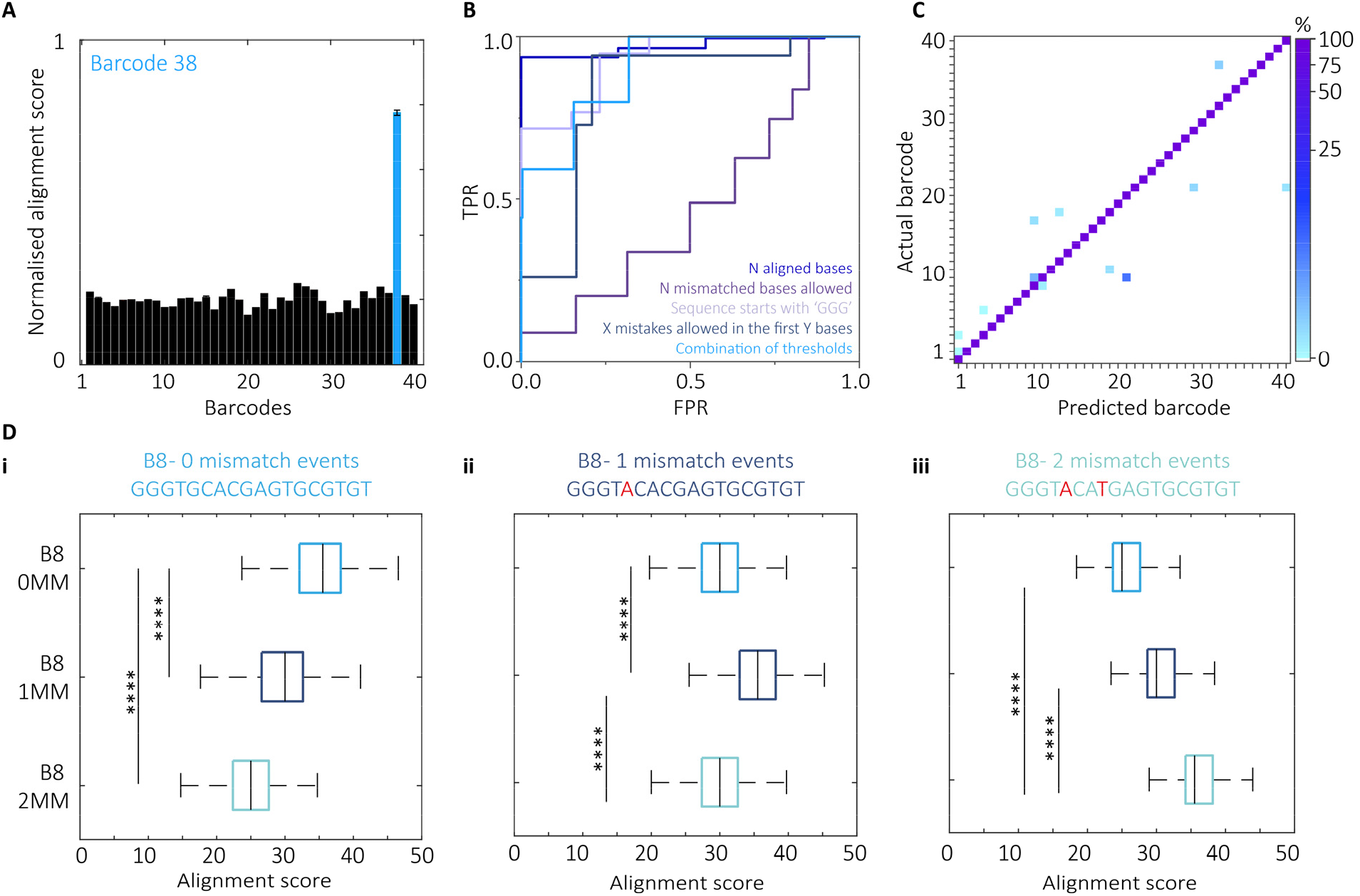
Highly accurate detection of nucleic acid barcodes. (a) Alignment scores of a barcoded probe 38 experiment. Sequenced events were aligned against a barcode library containing all 40 barcoded probe sequences. The alignment score for barcoded probe 38 was significantly higher (alpha = 0.0001, ANOVA, N=3, n=168-1845) than all other alignment scores observed. (b) ROC showing sensitivity and specificity of various alignment thresholds including N aligned bases; N mismatched bases allowed; sequence starts with *X; X* mismatches allowed in first *Y* bases and a combination (n=3). (c) Confusion matrix of alignment accuracy of single barcoded probe experiments. Accuracy was >95 % for all 40 barcoded probes tested (N=3, n=15489). (d) Alignment score of barcoded probe sequences with 1 and 2 mismatched bases. A multiplexed experiment containing three barcoded probes (B8, B8-1MM, B8-2MM) shows a significant difference between alignment scores (alpha = 0.0001, ANOVA, N=3, n=44015). (d,i) B8 events had the highest alignment score for the B8 sequence (d,ii) B8-1MM events had the highest alignment score for the B8-1MM sequence (d,iii) B8-2MM events had the highest alignment score for the B8-2MM sequence.

### Hybridisation of barcoded probes with miRNA

Many miRNAs share similar sequences.^51^ To demonstrate sequence-specific miRNA detection, a mixture of 40 barcoded probes was incubated with each miRNA individually (Fig. 3A). It was found that the percentage delayed events of each barcoded probe increased significantly (p <0.01) when in the presence of its corresponding target miRNA. We assessed the increase in percentage delayed events by comparing the miRNA sequence homologies (Supplementary Fig. 3a). Using a threshold of ≥90% miRNA sequence similarity and a significant increase in percentage delay (p≤0.01), we found that 2.65% of all classifications were true positive (TP), 0.95% false positive (FP), 0% false negative (FN) and 96.40% true negative (TN) (Fig. 3b), resulting in a platform accuracy of 99.05%, specificity of 99.03%, and sensitivity of 100%. Importantly, we observed that the barcode sequence does not influence the percentage delayed events detected for our probes (Supplementary Fig. 3b). In single barcoded probe experiments (e.g. barcoded probe 38 and miR-221-5p), the total event time increased when 50 nM of miRNA was present compared to the control (Fig. 3c and Supplementary Fig. 2b). However, it is known that the DNA-controlling motor enzyme does not move the probe unidirectionally through the pore; therefore, the speed at which the barcoded probes are translocated varies^52^. Hence, rather than relying on the translocation time, the moving standard deviation of the electrical signal was used to identify signal delays. Using this method, it was possible to determine a concentration-percentage delay curve for each barcoded probe and miRNA combination (R^2^ ≥ 0.989) (Fig. 3s, Supplementary Fig. 4).

**Fig. 3.**
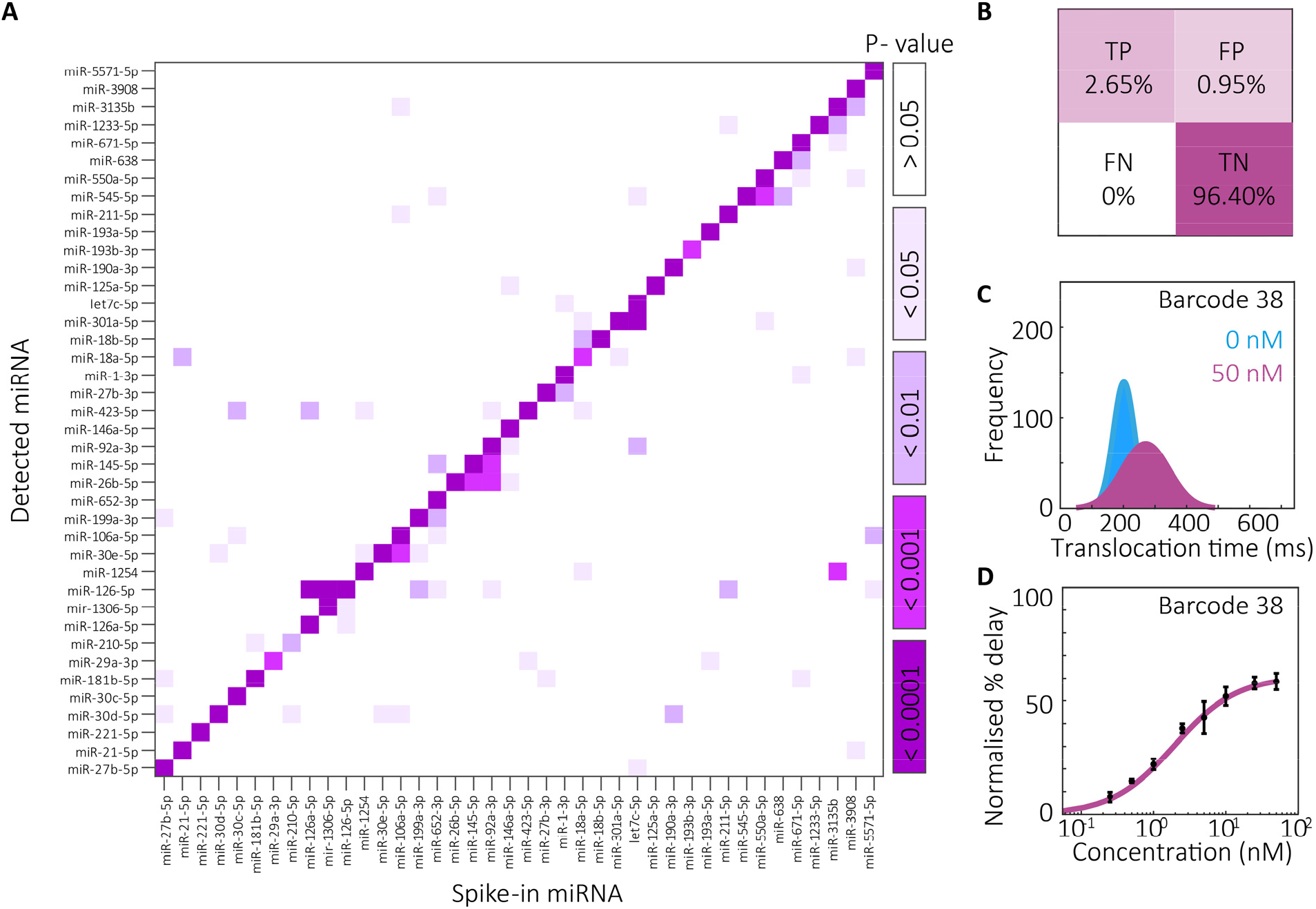
Sensitivity and selectivity of single miRNA detection. (a) Heatmap showing significance for selective miRNA binding. 40 barcoded probes were incubated with each miRNA, respectively. A significant increase (p≤ 0.01) in percentage delayed events was observed for all barcoded probes when in the presence of its miRNA target (N=3, n=99-1046). (b) Confusion matrix showing true positive, false positive, false negative and true negative occurrence. The accuracy was 99.05%, specificity was 99.03%, and sensitivity was 100%. (c) Translocation time of single barcoded probe and single miRNA experiments (N=3, n=3610,3904). (d) Concentration-percentage delay relationship of single miRNA and single barcoded probe experiment (barcoded probe 38, miR-221-5p). Data were fitted using the Michaelis Menten equation (K_d_= 1.77 nM and V_max_= 61.03 %, N=5, n=26798-308954).

### Multiplexed detection of miRNAs, proteins, and small molecules

Barcoded probes were incubated with synthetic miRNAs to identify if event delays could be observed in multiplexed conditions (Fig. 4a). It was found that the mean percentage delayed events increased when miRNAs were present (0 nM vs 50 nM 12.32 ±0.93% vs 54.84 ±2.41%, Student’s t-test p≤0.0001, n= 5) (Fig. 4B, i). Without demultiplexing, it was found that there was a linear increase in percentage delayed events between 0.1 nM and 10 nM (R^2^ ≥ 0.988) followed by a plateau (Fig. 4B, ii). The signal was demultiplexed using our alignment protocol, identifying individual populations of each barcoded probe. We then constructed bespoke concentration-percentage delay curves for each probe (Fig. 4b, iii). We next tested our ability to quantify multiple miRNAs using a single-blinded test. 40 barcoded probes were incubated with a mix of 40 miRNAs at various concentrations, ranging from 0.5 nM to 20 nM (Fig. 4c). The event files were sent to a blinded researcher who determined the percentage delay of each barcoded probe and interpolated the value with the standard curves to determine the concentration of the miRNAs. After estimation of each miRNA concentration, the experiment was unblinded and compared to the actual concentration (Fig. 4c, i and Supplementary Fig. 4). A residual analysis was performed of the data, where the percentage difference between the predicted and actual concentration was calculated (Fig. 4C, ii), all predictions were within one order of magnitude of the true concentration.

**Fig. 4.**
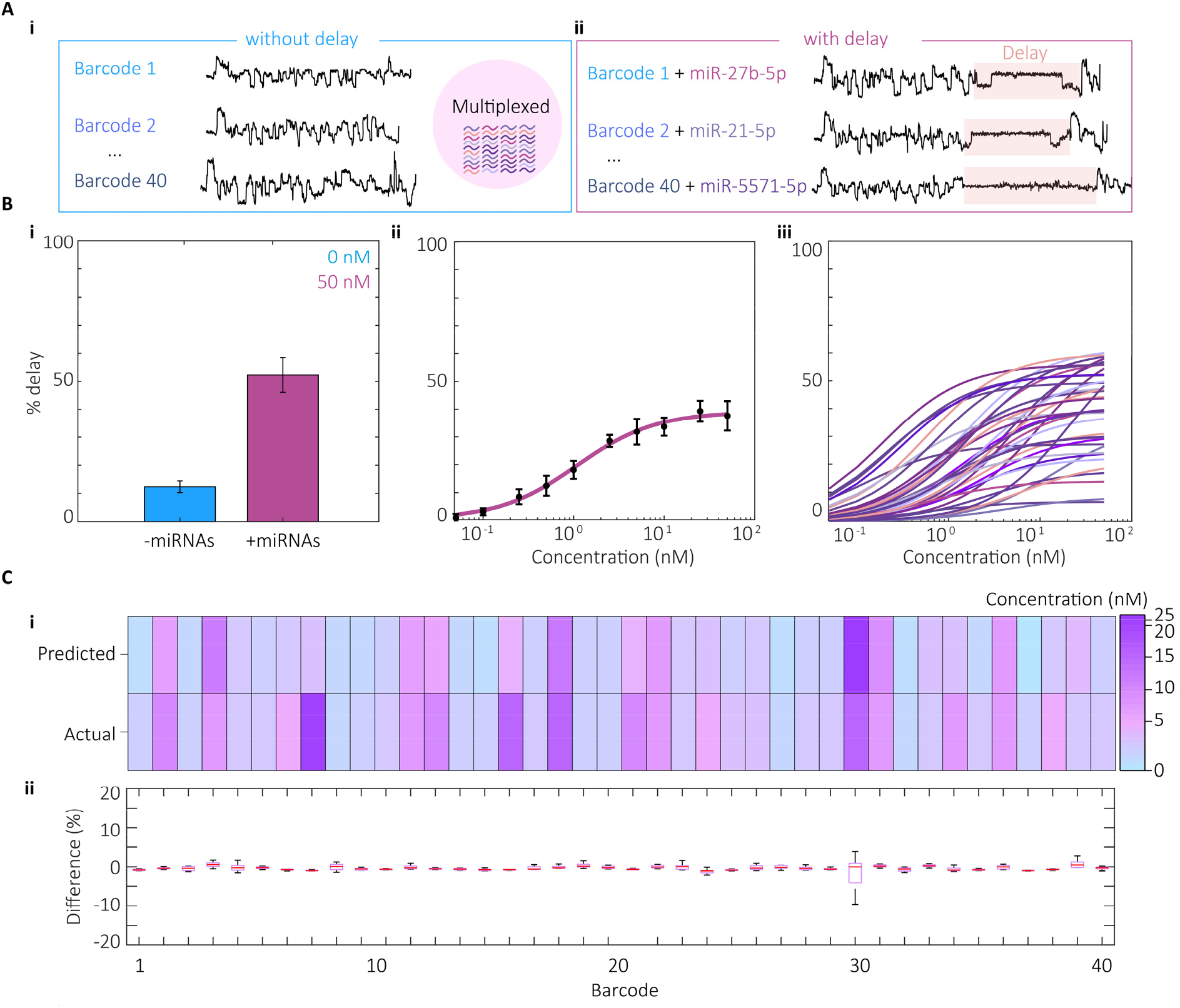
Multiplexed detection and quantification of miRNAs. (a, i) Example current traces for barcodes without delay and (ii) with delay. (b, i) Mean percentage delay +/- 50 nM miRNAs (****p≤0.0001, N=5, n=76841,69983). (b, ii) Multiplexed concentration-percentage delay curves of 40 barcoded probes and the corresponding 40 miRNAs. A curve was fitted using the Michaelis Menten equation (K_d_ = 1.09 nM, V_max_ = 38.69%, N=5, n=26798-308954). (b, iii) Concentration-percentage delay curves of 40 individual barcoded probes derived from a multiplexed experiment. Curves were fitted with Michaelis Menten equation (N=5, n=26798-308954). (c) Single-blinded prediction of miRNA concentration and comparison to known concentration. (c, i) Heatmap of concentrations (N=4, n=6189-71031). (c, ii) Residual analysis of miRNA predictions (N=4, n=6189-71031).

Multiplexed detection of analytes of different molecular species has not yet been realised. Due to its adaptability, this platform has the potential to detect proteins, miRNAs and small molecules in a single experiment. Barcoded probes were designed with aptamers specific for each protein and small molecule, (Supplementary Table 3). In single barcoded probe experiments, a significant increase in translocation time could be observed when in the presence of its corresponding protein: Thrombin (Fig. 5a, Student’s t-test, *p≤0.05); B-type natriuretic peptide (Fig. 5b, Student’s t-test, **p≤0.01); cardiac troponin-T (Fig. 5c, Student’s t-test, ****p≤0.0001) and cardiac troponin-I (Fig. 5d, Student’s t-test, ***p≤0.001). This corresponded with a significant increase in the percentage delayed events (Fig. 5a-d), suggesting that the detection of proteins is possible with this method. Moreover, a significant increase in translocation time and percentage delayed events was observed when serotonin (150 nM) was incubated with its corresponding barcoded probe (Fig. 5E, Student’s t-test, *p≤0.05).

**Fig. 5.**
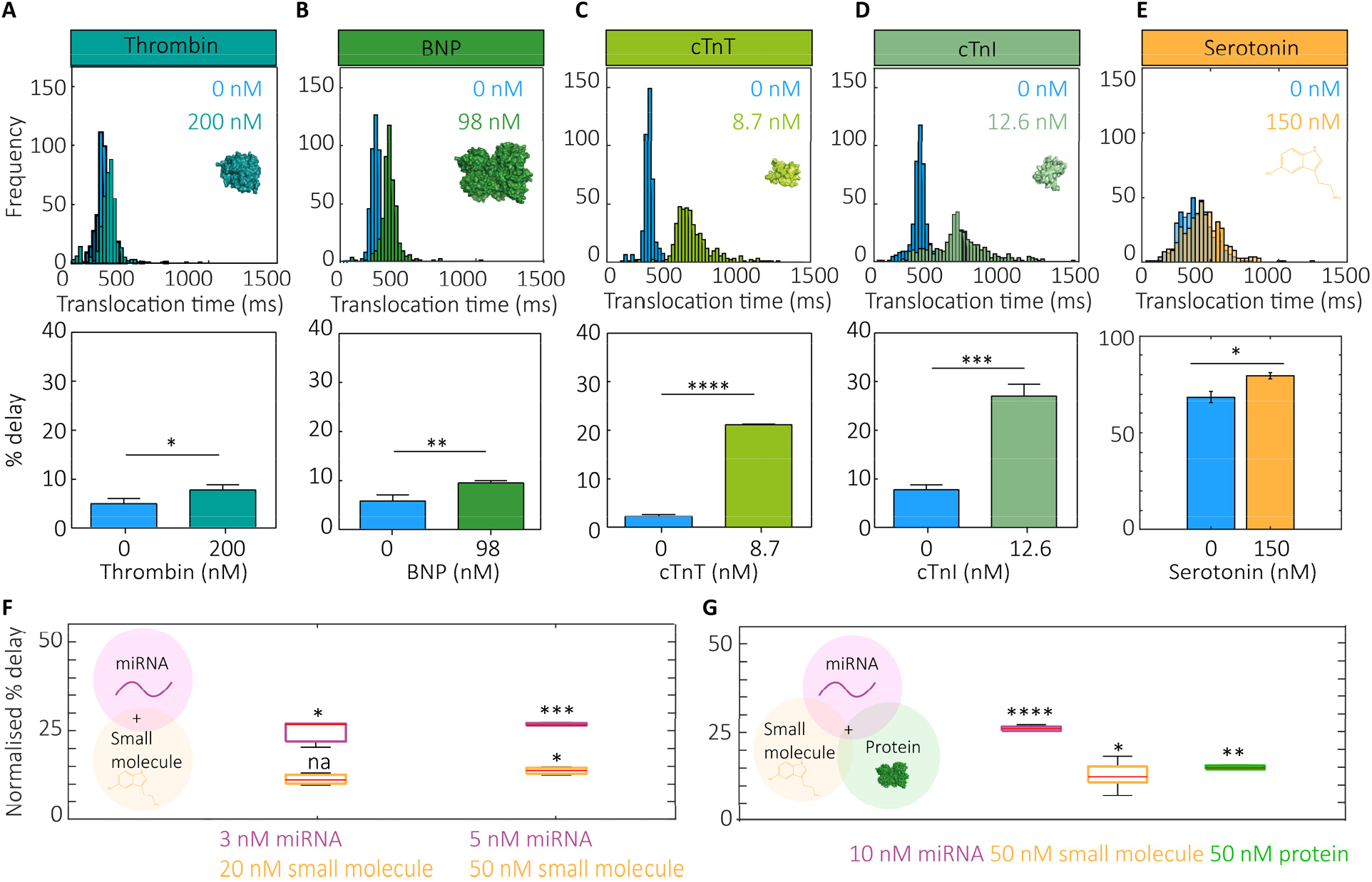
Protein and small molecule detection. Translocation time and percentage delayed events of barcoded probes +/-each corresponding protein or small molecule. (a) Thrombin, (b) BNP, (c) cTnT, (d) cTnI, and (e) serotonin. Significance was determined by Student’s t-test (*p≤0.05, **p≤0.01, ***p≤0.001, ****p≤0.0001, N=3, n=1024-3395). (f) Multiplexed small molecule and miRNA detection. Normalised percentage delay of barcoded probes is shown. Significance was determined by Student’s t-test (*p≤0.05, ***p≤0.001, N=3, n=1307-1745). (g) Multiplexed small molecule, miRNA, and protein detection. Normalised percentage delay of barcoded probes is shown. Significance was determined by Student’s t-test (*p≤0.05, **p≤0.01, ***p≤0.001, N=3, n=667-1323).

Multiplexed experiments were conducted by observing analytes of different molecular species. Firstly, a duplex experiment of a small molecule and a miRNA was conducted. Incubation of barcoded probes with 50 nM small molecule and ≥3 nM miRNA significantly increased the percentage delayed events for each probe (Fig. 5f). We also performed a triplex experiment observing a miRNA, small molecule, and protein simultaneously. The percentage delayed events for each barcoded probe increased significantly from the controls when miRNA (10 nM), small molecule (50 nM) and protein (50 nM) were present (Fig. 5g).

### Detection of miRNAs in human serum

Blood serum from eight healthy participants was tested for the presence of 40 miRNAs and compared to the negative control (no serum, only buffer). The list of miRNAs selected for detection in this assay have all been previously associated with cardiovascular disease (Supplementary Table 1). The mean percentage delay of all miRNAs (not demultiplexed) was increased in each of the 8 participant samples in three independent experiments, however no significance was determined (Fig. 6a). Interestingly, when the signal was demultiplexed, significant changes (Student’s t-test, p≤0.05) in percentage delay of miRNAs could be observed (Fig. 6b). Of the 40 miRNAs tested, we observed a significant increase in 24 miRNAs across all 8 participants (Fig. 6b and c, Supplementary Fig. 7). Each participant had an average of 3 elevated miRNAs (minimum= 1, maximum= 5). The most frequently observed miRNAs across all samples were: miR-1233-5p, miR146a-5p, miR211-5p, miR30c-5p, miR18a-5p, miR-126-5p and miR193b-3p.

**Fig. 6.**
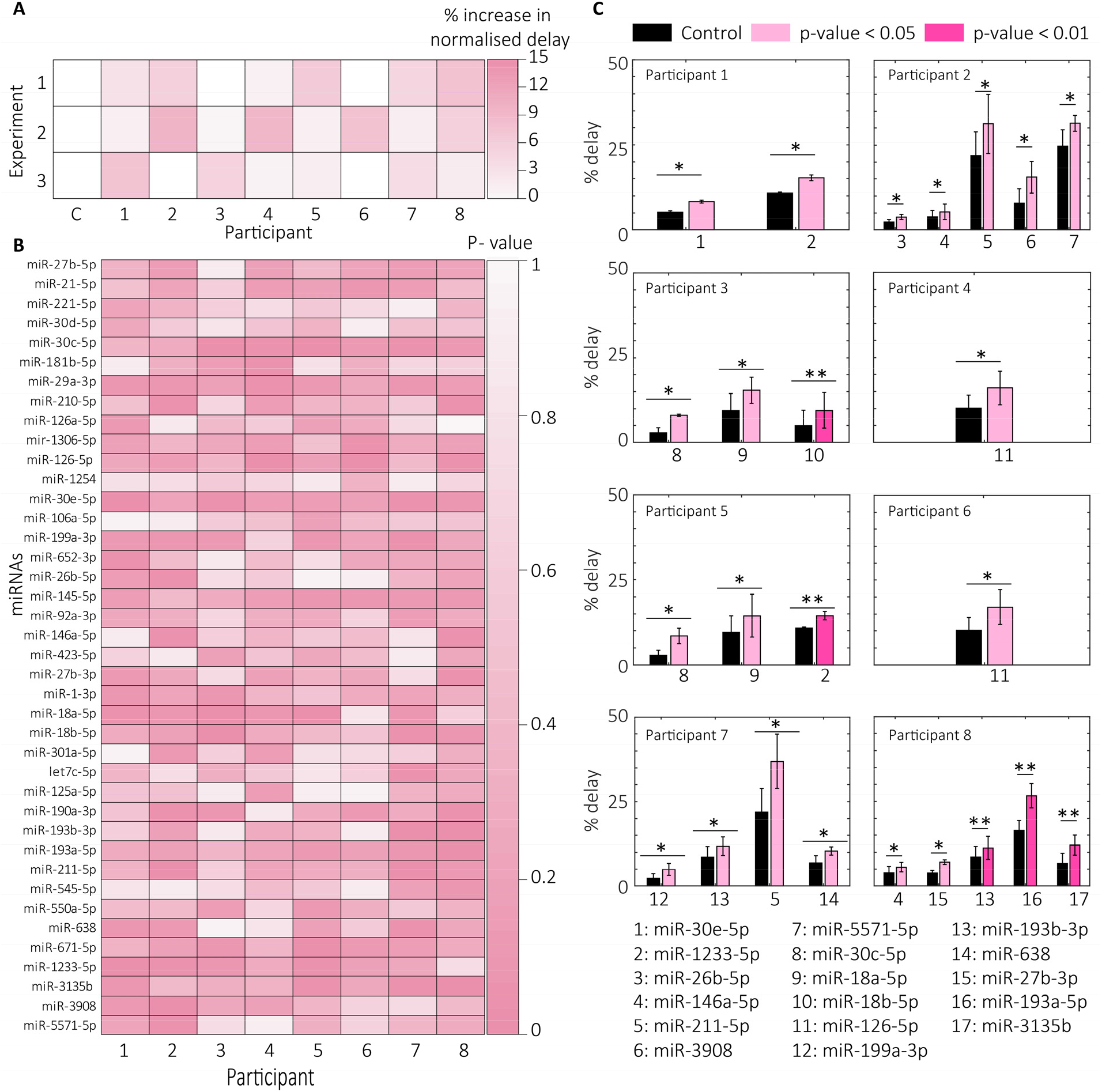
Multiplexed detection of miRNAs in human serum. (a) Mean percentage delay of barcoded probes (not demultiplexed). Serum was incubated with 40 barcoded probes and compared to zero serum control (N=3, n=17930-25776) (b) Heatmap of p-values comparing percentage delayed events +/− serum (Student’s t-test, N=3, n=17930-25776). (c) Significantly upregulated miRNAs per patient. Significance determined by Student’s t-test (*p≤0.05, **p≤0.01).

## Conclusions

There is a genuine clinical need to develop platforms which can rapidly detect multiple biomarkers in patient samples. Through expanding the panel of molecules observed, it is possible that disease subpopulations could be identified and treatment regimens optimised. Detection of biomarkers of different molecular species within the same sample could also massively reduce the analysis pipeline.

Our data shows that barcoded probes, in combination with the ONT sequencing platform, offer a highly accurate and sensitive analyte sensing platform with single-molecule resolution. We were able to detect 40 different barcoded probes in the same sample and construct independent concentration-delay curves. We then used these standard curves to predict miRNA concentrations in a single-blinded experiment. The multiplexed method was then applied to multiple molecular species allowing us to observe an increase in percentage delayed events due to proteins and small molecules (as well as miRNAs). Finally, we showed that the platform is compatible with human serum, indicating its potential as a multiplexed biomarker detection platform.

In the platform’s current configuration, it is possible for the assay time to be less than 60 minutes. This is a significant improvement compared to other technologies, which take several hours or even days to complete^53^. Moreover, since the MinION device is highly portable, assays could be performed away from the clinic. The speed and portability of the platform offer further advantages over other biomarker detection strategies. In principle, this technology could also be adapted to allow for the pooling of samples from multiple patients, lowering the assay cost and time.

The data presented have shown that it is possible to multiplex the detection of at least 40 analytes. The platform can correctly distinguish between barcodes with a single nucleotide change in the sequence. Interaction of probes was predicted (Supplementary Fig. 5); however, there was no clear relationship between dimer formation and event capture rate. This indicates that any future applications of this technology will not be limited by barcode sequences but perhaps the selectivity of the binding regions. Our data suggest that miRNA sequences with a similarity ≥90% have the potential to hybridise with the incorrect probe (Supplementary Fig. 7); strategies to distinguish between such miRNAs must therefore be further developed. There is also potential for the large number of biomarkers assessed to result in “data blindness” to address this, a graphical interface or risk scoring method must be developed to assist the end user. When translating this device to the clinic, much work is required to deliver information to clinicians in a straightforward manner which can help inform decisions. This could likely be determined through the application of neural networks or AI systems to spot the patient sub-population “fingerprints”.

Currently, the detection limit of the platform for miRNA experiments is c.50 pM. There is potential to push the technology further by reducing the false negative rate of event alignment. At the moment, there are a large number of events which are rejected due to the thresholds applied to ensure high accuracy. These events are not considered for delay analysis since we are unable to distinguish them from any false positives. One way to reduce this event loss and increase events available for delay analysis would be to switch to a different nanopore model.

A particular issue of assaying blood serum is that nanopore lumen are small and are easily blocked by proteins. Pore blockage reduces the barcoded probe capture rate, meaning assay times must be increased or repeated. To reduce the blockage of nanopores by proteins in this study, we added a filtration step which removed the largest serum proteins (>10 kDa). In future applications, it would be preferable to develop our methodology further to reduce the sample processing required before detection whilst also reducing the unwanted pore-blocking.

The platform shows great potential for use in clinical environments, for example, to offer expansive, longitudinal disease tracking or early disease detection. With further optimisation, this strategy could significantly reduce testing time, assay cost and sample volume whilst increasing the data available to the clinician.

## Materials and Methods

### Chemicals and Materials

All flow cells (MinION and Flongle) were provided by ONT, UK. All probe sequences were custom designed. Barcoded probes and miRNAs were synthesised by Integrated DNA Technologies, USA (Tables 1 and 3). Proteins were obtained from the following: Thrombin (RP-43100, Invitrogen, USA); B-type natriuretic peptide (4095916, Bachem, Switzerland); cardiac troponin I (Z03320, Genscript Biotech, USA); cardiac troponin T (230-00048, Ray Biotech, USA). Small molecule: Serotonin (H9523, Sigma-Aldrich, UK). Ampure XP magnetic beads (for purification of DNA barcoded probes) were purchased from Beckman Coulter, USA. TA ligase was acquired from New England Biolabs, USA. All chemicals used for buffer preparation were obtained from Sigma-Aldrich (UK), Roche, Switzerland or VWR Chemicals, USA. In all experiments, DNA lo-bind tubes were used (Eppendorf, Germany).

### Human donor blood serum

Human samples used in this research project were obtained from the Imperial College Healthcare Tissue Bank (ICHTB). ICHTB is supported by the National Institute for Health Research (NIHR) Biomedical Research Centre based at Imperial College Healthcare NHS Trust and Imperial College London. ICHTB is approved by Wales REC3 to release human material for research (22/WA/0214), and the samples for this project (R22016) were issued from subcollection reference number NHL_FN_021_028.

Venous blood was collected in red-topped vacutainers (Beckton Dickinson, USA) and allowed to clot at room temperature for 15 min before centrifugation at 3000 x g, 15 min, 4 ^o^C. The resulting serum was then aliquoted into small volumes and frozen at −80 ^o^C until use. The serum was filtered using a 10 kDa molecular weight cut-off spin filter (Sartorius, Germany) before incubation with barcoded probes.

### Barcoded probe design

The complete carrier design consists of 3 sections named ‘adapter’, ‘barcode region’ and ‘target binding region’ (Fig. 1A); when fully assembled, this was called a ‘barcoded probe’. The adapter section is an ONT Y-adapter; it consists of; i) a leader (which facilitates threading into the nanopore); ii) a tether (to enhance the capture rate), and iii) a motor protein (to control the translocation of the barcoded probe through a nanopore). The barcode region consists of a polynucleotide identifier and is followed by spacer nucleotides (to separate the barcode and target binding regions). The target binding region consists of a DNA aptamer or complementary miRNA sequence, depending upon the species of the target analyte.

### Barcoded probe preparation

Each probe was incubated with ligation c-strand in a molar ratio of 1:3 in nuclease-free water at room temperature for 1 h. The resultant mix was combined with 10 nM adapter and an equal volume of TA ligase master mix (New England Biolabs). The mix was centrifuged at 4°C for 1 min and then incubated at room temperature for 20 min. Probes were purified using the SPRI (solid phase reversible immobilisation) method. Ampure XP beads (Beckman Coulter, USA) were added at 1.4 times the total solution volume. The beads (with probes bound) were washed two times with a short fragment buffer (ONT, UK). After the washes, the beads were resuspended in nuclease-free water, causing the probes to be released. 100 nM tether was then added to the probes along with sequencing buffer. The 2x sequencing buffer contained: 700 mM KCl, 50 mM HEPES, 100 mM MgCl_2_, 100 mM ATP, 4.4 mM EDTA (pH= 8.0). Barcoded probes were incubated with the target analyte for 30 min at room temperature before loading into flow cells.

Pre-prepared mixes of barcoded probes are resilient to multiple freeze-thaw cycles and long-term storage at −20°C (Supplementary Fig. 8). Hybridisation dynamics of miRNAs with barcoded probes were also investigated, revealing that incubation of 5-10 minutes is sufficient to reach equilibrium of probe-analyte binding (Supplementary Fig. 9). This can potentially reduce the sample preparation time and assay variability significantly.

### Sequencing experiments and data acquisition

All sequencing experiments were performed at 34°C using either the MinION or Flongle sequencing device (ONT, UK). The MinION/ Flongle was connected through a USB 3.0 port to a PC with a minimum of 16 GB RAM. A membrane check was performed before each run to determine the integrity of the membrane and to identify how many nanopores were active. Before each experiment, flow cells were flushed with 2 x 500 μL (MinION) or 1 x 150 μL (Flongle) sequencing buffer. The volume for each sequencing experiment was determined by the flow cell used; 150μL (MinION) or 30μL (Flongle). After each experiment, the MinION flow cell was washed with the flow cell wash kit (ONT, UK) according to the manufacturer’s protocol.

Data collection was performed using the proprietary software MinKNOW (ONT, UK). Basecalling was either performed in real-time (MinKNOW) or offline within a custom-written MATLAB script, ‘The Nanopore app’, previously published by our group^48^.

### Event analysis

Barcoded probe translocations were identified and analysed using the following workflow: 1) Event identification; 2) Event basecalling; 3) Event alignment; 4) Event delay analysis.

Event identification included the tracking, and then subtraction of the baseline signal. A cut-off threshold was then determined based on the background noise (30-40 standard deviations above mean noise level). ‘peakfinder’ function in MATLAB was used to spot events. Any events identified that were shorter than 100 ms were excluded at this stage.

Event basecalling was performed with the Guppy basecaller using the high accuracy (HAC) flip-flop model (ONT, UK).

Each basecalled event was aligned against a library of barcode sequences. Each event was attributed to one of the library sequences based on an alignment scoring method. Further thresholds were applied to ensure that only true positive barcode events were retained for further analysis. These thresholds were: i) sequence starts with “GGG”, ii) ≥15 bases aligned to the library sequence, iii) 1 mismatched base in the first 10 bases, and iv) ≤5 mismatched bases in the entire sequence.

In order to distinguish between analyte-bound and unbound barcoded probe events, we performed analysis of each event. When an analyte is bound to the barcoded probe, there is a ‘quiet’ sojourn in the electrical signal, which persists until the analyte is dissociated from the probe, at which point it can complete its translocation. An event is defined as delayed if the moving standard deviation of the signal is less than the threshold of 0.003 for a period greater than 10 bins (each event signal is separated into a total of 75 bins). All other events are defined as having no delay.

## Data availability

The data that support the plots within this paper and other findings of this study are available from the corresponding author upon reasonable request. Source data for the figures and supplementary figures are provided as a Source Data file.

## Code availability

The Nanopore App used to analyse the data is available from the corresponding author upon reasonable request.

## Acknowledgements

We thank Miruna Cretu for helpful discussion on data analysis. We thank Dr Liang Xue for his involvement in early stages of this project and contributing to helpful discussions. C.K. acknowledges studentship funding from Oxford Nanopore Technologies and the Institute of Chemical Biology at Imperial College London. A.P.I. and J.B.E. acknowledge support from BBSRC grant BB/R022429/1, EPSRC grant EP/V049070/1, and Analytical Chemistry Trust Fund grant 600322/05. This project has also received funding from the European Research Council (ERC) under the European Union’s Horizon 2020 research and innovation programme (grant agreements No 724300 and 875525).

## Contributions

C.K., B.R., A.P.I. and J.B.E. conceived and designed experiments. C.K. and B.R. performed experiments and wrote the manuscript. J.B.E wrote and developed the NanoporeApp analysis software. C.L. collected and prepared of human serum samples. B.R., F.S.N. and J.G. obtained the ethical approval for the collection and storage of human serum samples. R.G., J.G., A.P.I, and J.B.E. were responsible for the conception and supervision of the project. All authors edited the manuscript.

