## Supplementary Information for "Highly multiplexed detection of microRNAs, proteins and small molecules using barcoded molecular probes and nanopore sequencing"

Supplementary table 1: Table of barcoded probe sequences. All probes were purchased from IDT with high purity and desalting. iSpC3= C3 Spacer phosphoramidite. Associated cardiac diseases: ACS Acute Coronary syndrome; AMI Acute Myocardial Infarction; AS Aortic Stenosis; ASVD Atherosclerosis; CAD Coronary Artery Disease; CF Cardiac Fibrosis; DCM Dilated cardiomyopathy; HF Heart failure; HFpEF Heart failure with preserved ejection fraction; HFrEF Heart failure with reduced ejection fraction; Hyp Hypertension; NSTEMI Non-ST-elevated myocardial infarction; SCD Sudden cardiac death; STEMI ST-elevated myocardial infarction. $K_d$ *and* $V_{max}$ *values* were obtained by fitting multiplexed experiments with the Michaelis Menten equation.

| Number | Barcode name | Target sequence (miRNA sequence) | Associated cardiac disease | Binding sequence | Barcode sequence | Multiplexed $K_d$ (nM) | Multiplexed $V_{max}$ |
|---|---|---|---|---|---|---|---|
| 1 | Adapter_barcode_c-hsa-miR-27b-5p | CAAGTGGTTAGTCGATTCGAGA | CF / Hyp / ACS / HF | GTTCACCAATCAGCTAAGCTCT | GGGATCGCTACGCCTTCGGCTCGTAATCATAGTCGAGT | 0.976 | 48.445 |
| 2 | Adapter_barcode_c-hsa-miR-21-5p | AGTTGTAGTCAGACTATTCGAT | NSTEMI / ASVD | TCAACATCAGTCTGATAAGCTA | GGGAGCTCAGAGCAGGTCACTCAAGATACGAGCTGCGT | 4.172 | 62.005 |
| 3 | Adapter_barcode_c-hsa-miR-221-5p | TTTAGATGTAACATACGGTCCA | SCD | AAATCTACATTGTATGCCAGGT | GGGGTAAGTCTGCATCAGCGCGCGGCTGTGCGAGGATA | 2.107 | 32.552 |
| 4 | Adapter_barcode_c-hsa-miR-30d-5p | GAAGGTCAGCCCCTACAAATGT | STEMI | CTTCCAGTCGGGGATGTTTACA | GGGCTACGACAGTACGCTAGCAAGGATAGACACTACGA | 9.060 | 64.651 |
| 5 | Adapter_barcode_c-hsa-miR-30c-5p | CGACTCTCACATCCTACAAATGT | CF / HF | GCTGAGAGTGTAGGATGTTTACA | GGGTACTGAACACAAGTTCGTCGTCGAGCAATCACAAT | 14.363 | 34.463 |
| 6 | Adapter_barcode_c-hsa-miR-181b-5p | TGGGTGGCTGTCGTTACTTACAA | AS | ACCCACCGACAGCAATGAATGTT | GGGCTAGTGCGCAGTTGTCTCGGCGGAGTTGAGACTGA | 1.453 | 62.109 |
| 7 | Adapter_barcode_c-hsa-miR-29a-3p | ATTGGCTAAAGTCTACCACGAT | CF / HF/ STEMI | TAACCGATTTCAGATGGTGCTA | GGGGATCATGGTAGTCTTCAAGATCGAGTATGTCTGTC | 1.881 | 24.849 |
| 8 | Adapter_barcode_c-hsa-miR-210-5p | GTCACACGCCACCCGTCCCCGA | HF | CAGTGTGCGGTGGGCAGGGGCT | GGGGTTCACATCAAGGTCATACCGCGAGTTCTATTTTA | 1.407 | 22.671 |
| 9 | Adapter_barcode_c-hsa-miR-126-5p | GCGCATGGTTTTCATTATTAC | HF | CGCGTACCAAAAGTAATAATG | GGGGCTTGGGGGATAGATGTGCCCCGCGCATCGGACCT | 0.832 | 47.446 |
| 10 | Adapter_barcode_c-hsa-mir-1306-5p | ACCTGCAAACGTCCCCTCCACC | AMI | TGGACGTTTGCAGGGGAGGTGG | GGGATGACACACGTTTTCGATAGGGACGCCGACTTTAA | 0.126 | 27.487 |
| 11 | Adapter_barcode_c-hsa-miR-126-5p | GCGCATGGTTTTCATTATTAC | HF | CGCGTACCAAAAGTAATAATG | GGGTGATAATAAGACCTGACAGACAATAGGGAGAACTC | 0.193 | 56.250 |
| 12 | Adapter_barcode_c-hsa-miR-1254 | TGACGTCCGAGGTCGAAGGTCCGA | HF | ACTGCAGGCTCCAGCTTCCAGGCT | GGGTTAGTAATCAAGTCTGATCGTAATAGCTAAGTCAT | 1.491 | 57.875 |
| 13 | Adapter_barcode_c-hsa-miR-30e-5p | GAAGGTCAGTTCCTACAAATGT | DCM / AS | CTTCCAGTCAAGGATGTTTACA | GGGCCTGTTCTAATTGCGCGGAGAGGCGAGATGTTTCT | 2.244 | 41.573 |
| 14 | Adapter_barcode_c-hsa-miR-106a-5p | GATGGACGTGACATTCGTGAAAA | DCM | CTACCTGCACTGTAAGCACTTTT | GGGTGGGATACTGACGTGCCGGAATACCCAGACGTGCC | 2.451 | 30.826 |
| 15 | Adapter_barcode_c-hsa-miR-199a-3p | ATTGGTTACACGTCTGATGACA | AMI / HF | TAACCAATGTGCAGACTACTGT | GGGTGATCGGCACCTAAACGCATTAGCCCTGCAATACG | 1.443 | 41.053 |
| 16 | Adapter_barcode_c-hsa-miR-652-3p | GTGTTGGGATCACCGCGGTAA | ACS | CACAACCCTAGTGGCGCCATT | GGGAGCTGCTCGGAAGCCATAAGGTACTTTAATTTGGG | 2.682 | 18.111 |
| 17 | Adapter_barcode_c-hsa-miR-26b-5p | TGGATAGGACTTAATGAACTT | AS | ACCTATCCTGAATTACTTGAA | GGGCACGGATTTCTATATTGCTCAACCAGGCAGCGCAA | 0.216 | 49.819 |
| 18 | Adapter_barcode_c-hsa-miR-145-5p | TCCCTAAGGACCCTTTTGACCTG | ACS / CAD / STEMI | AGGGATTCCTGGGAAAACTGGAC | GGGATTGTGCGCTTTTCTCCATGCGTCTTAGACATCTC | 5.260 | 64.427 |
| 19 | Adapter_barcode_c-hsa-miR-92a-3p | TGTCCGGCCCTGTTCACGTTAT | ACS | ACAGGCCGGGACAAGTGCAATA | GGGGTGAAATCCCCGTCTAGGTTATGGCTGGGGGGATT | 0.866 | 30.393 |
| 20 | Adapter_barcode_c-hsa-miR-146a-5p | TTGGGTACCTTAAGTCAAGAGT | ACS / AMI / STEMI / HF | AACCCATGGAATTCAGTTCTCA | GGGCGTAAACTTATCACGACACAATGAACAAGCCTGCA | 0.568 | 60.163 |
| 21 | Adapter_barcode_c-hsa-miR-423-5p | TTTCAGAGCGAGAGACGGGGAGT | HF | AAAGTCTCGCTCTCTGCCCCTCA | GGGGGCAGTGTCCGAGCGTCCTCAATCATGAGCGATTC | 2.041 | 38.207 |
| 22 | Adapter_barcode_c-hsa-miR-27b-3p | CGTCTTGAATCGGTGACACTT | AS | GCAGAACTTAGCCACTGTGAA | GGGCGTCCAGACTTAATGTCTGCTCACTGACATCGCGA | 2.468 | 52.598 |
| 23 | Adapter_barcode_c-hsa-miR-1-3p | TATGTATGAAGAAATGTAAGGT | NSTEMI /ACS | ATACATACTTCTTTACATTCCA | GGGATTAGCGGAACCAAACCCAGGAAGGCTTGAAGGCG | 0.626 | 14.264 |
| 24 | Adapter_barcode_c-hsa-miR-18a-5p | GATAGACGTGATCTACGTGGAAT | CAD | CTATCTGCACTAGATGCACCTTA | GGGAACCTTAGGGGCCTCGAATCTTTGAGACGACTAGG | 0.587 | 6.944 |
| 25 | Adapter_barcode_c-hsa-miR-18b-5p | GATTGACGTGATCTACGTGGAAT | HF | CTAACTGCACTAGATGCACCTTA | GGGTAATTACTGCCCCACCATGACATTTTAATAGCAGT | 0.992 | 30.556 |
| 26 | Adapter_barcode_c-hsa-miR-301a-5p | TCATCACGTTATTTCAGTCTCG | HF | AGTAGTGCAATAAAGTCAGAGC | GGGGACCTTGAGACAGAACTTATCAATGTACAACTGAA | 1.081 | 30.850 |
| 27 | Adapter_barcode_c-hsa-let7c-5p | TTGGTATGTTGGATGATGGAGT | CAD | AACCATACAACCTACTACCTCA | GGGCGAAGGATTGGCCCCCCGATTACCACCGCCGTGAG | 0.306 | 39.098 |
| 28 | Adapter_barcode_c-hsa-miR-125a-5p | AGTGTCCAATTTCCCAGAGTCCCT | HFrEF | TCACAGGTTAAAGGGTCTCAGGGA | GGGAATTGCCAACAGGTCAAGCCCTGTTCTCACTGGTC | 0.251 | 52.617 |
| 29 | Adapter_barcode_c-hsa-miR-190a-3p | TCCTTATACAAACTATATATC | HFpEF | AGGAATATGTTTGATATATAG | GGGCAGTGCTTGCCCCCAGTAGAGTGTGGAAGGGCATA | 0.692 | 24.663 |
| 30 | Adapter_barcode_c-hsa-miR-193b-3p | TCGCCCTGAAACTCCCGGTCAA | HFrEF /STEMI | AGCGGGACTTTGAGGGCCAGTT | GGGAAAGCCCACTCTCCACACTTCAAGGTTAAATGGCG | 6.454 | 21.231 |
| 31 | Adapter_barcode_c-hsa-miR-193a-5p | AGTAGAGCGGGCGTTTCTGGGT | HF | TCATCTCGCCCGCAAAGACCCA | GGGGACTAATACAATCGGAAGCAACTCTCACGCCGCAC | 2.872 | 50.154 |
| 32 | Adapter_barcode_c-hsa-miR-211-5p | TCCGCTTCCTACTGTTTCCCTT | HFrEF / ASVD | AGGCGAAGGATGACAAAGGGAA | GGGCTACGATTCATGTCTCCCCCCACATATGATTGATC | 0.755 | 40.044 |
| 33 | Adapter_barcode_c-hsa-miR-545-5p | AGTAGATTATTTGTAAATGACT | HFpEF / STEMI | TCATCTAATAAACATTTACTGA | GGGTCGGGCGCTTAATCGCAATGTTCATCCGGAACGGA | 0.704 | 57.095 |
| 34 | Adapter_barcode_c-hsa-miR-550a-5p | CCCGAGAATGAGGGAGTCCGTGA | HFpEF / HF | GGGCTCTTACTCCCTCAGGCACT | GGGTAAATTCCACATTACGATAGCTGACGTCCTGGTAG | 4.246 | 8.531 |
| 35 | Adapter_barcode_c-hsa-miR-638 | TCCGGCGGTGGGCGGGCGCTAGGGA | HFrEF / HFpEF | AGGCCGCCACCCGCCCGCGATCCCT | GGGCACCCCCTAGCCTGACCTTAAATGATAATTCCTGT | 1.056 | 45.593 |
| 36 | Adapter_barcode_c-hsa-miR-671-5p | GAGGTCGGGGAGGTCCCGAAGGA | HFrEF | CTCCAGCCCCTCCAGGGCTTCCT | GGGTTTAGCCTAACCTCTGATGCATTGCACCGGAGTTC | 21.486 | 69.518 |
| 37 | Adapter_barcode_c-hsa-miR-1233-5p | ACGGCACGGGACCGGAGGGTGA | HFpEF | TGCCGTGCCCTGGCCTCCCACT | GGGCCCAATATACGCTGAACCTTCCATCCGATTTTCAG | 0.297 | 52.885 |
| 38 | Adapter_barcode_c-hsa-miR-3135b | GTGGTGACGTGAGCGAGGTCGG | HF | CACCACTGCACTCGCTCCAGCC | GGGCCTTGGAATTAGAACCGTGTGATTCTACGCCTAGG | 1.774 | 61.030 |
| 39 | Adapter_barcode_c-hsa-miR-3908 | TTTGTCAGATGGATGTAACGAG | HF | AAACAGTCTACCTACATTGCTC | GGGAGGACTTCTTCGGTGTAATCGGAGTACATCAATGT | 0.767 | 44.722 |
| 40 | Adapter_barcode_c-hsa-miR-5571-5p | CCCTCCGAGGAAACTCTTAAC | HF | GGGAGGCTCCTTTGAGAATTG | GGGAACGTACTTTGTGGGGATAAGCTGTACAGGGCTGT | 1.634 | 41.139 |

Supplementary table 2: Alignment of barcoded probe 38 sequenced events against barcoded probe 38 library sequence. The most common basecalled sequence, is shown in row two. The percentage value presents the occurrence of the most common base at that position in the alignment (n=30).

| | | | | | | | | | | | | | | | | | | | | | | | | | | | | | | | | | | | | | | |
|---|---|---|---|---|---|---|---|---|---|---|---|---|---|---|---|---|---|---|---|---|---|---|---|---|---|---|---|---|---|---|---|---|---|---|---|---|---|---|
| Library sequence Barcode 38 | G | G | G | C | C | T | T | G | G | A | A | T | T | A | G | A | A | C | C | G | T | G | T | G | A | T | T | C | T | A | C | G | C | C | T | A | G | G |
| Most common sequence | G | G | G | C | C | T | T | G | G | A | A | T | T | A | G | A | A | C | C | G | T | G | T | G | A | T | T | C | T | A | C | A | T | A | T | C | C | C |
| % | 100 | 100 | 100 | 100 | 100 | 99 | 100 | 99 | 99 | 98 | 99 | 98 | 99 | 95 | 93 | 95 | 97 | 96 | 96 | 91 | 91 | 90 | 91 | 90 | 90 | 91 | 93 | 77 | 50 | 40 | 34 | 46 | 33 | 31 | 34 | 32 | 41 | 41 |
| Sequenced event 1 | G | G | G | C | C | T | T | G | G | A | A | T | T | A | G | A | A | C | C | G | T | G | T | G | A | T | T | C | C | T | T | A | T | A | A | T | T | C |
| Sequenced event 2 | G | G | G | C | C | T | T | G | G | A | A | T | T | A | G | A | A | C | C | G | T | G | T | G | A | T | T | C | G | C | T | G | A | A | T | A | A | T |
| Sequenced event 3 | G | G | G | C | C | T | T | G | G | A | A | T | T | A | G | A | A | C | C | G | T | G | T | G | A | T | T | C | G | T | C | C | T | G | C | T | G | C |
| Sequenced event 4 | G | G | G | C | C | T | T | G | G | A | A | T | T | A | G | A | A | C | C | G | T | G | T | G | A | T | T | C | T | A | G | A | T | C | C | T | G | C |
| Sequenced event 5 | G | G | G | C | C | T | T | G | G | A | A | T | T | A | G | A | A | C | C | G | T | G | T | G | A | T | T | C | C | C | T | G | T | T | C | T | G | C |
| Sequenced event 6 | G | G | G | C | C | T | T | G | G | A | A | T | T | A | G | A | A | C | C | G | T | G | T | G | A | T | T | C | G | T | G | C | T | - | - | - | - | - |
| Sequenced event 7 | G | G | G | C | C | T | T | G | G | A | A | T | T | A | G | A | A | C | C | G | T | G | T | G | A | T | T | C | C | A | C | A | A | T | C | T | G | C |
| Sequenced event 8 | G | G | G | C | C | T | T | G | G | A | A | T | T | A | G | A | A | C | C | G | T | G | T | G | A | T | T | C | G | C | T | G | A | A | T | C | A | C |
| Sequenced event 9 | G | G | G | C | C | T | T | G | G | A | A | T | T | A | G | A | A | C | C | G | T | G | T | G | A | T | T | C | T | T | C | G | T | G | C | A | C | C |
| Sequenced event 10 | G | G | G | C | C | T | T | G | G | A | A | T | T | A | G | A | A | C | C | G | T | G | T | G | A | T | T | C | T | G | A | A | T | C | A | C | C | A |
| Sequenced event 11 | G | G | G | C | C | T | T | G | G | A | A | T | T | A | A | G | A | A | C | C | G | T | G | T | G | A | T | T | C | T | A | G | A | T | A | C | C | A |
| Sequenced event 12 | G | G | G | C | C | T | T | G | G | A | A | T | T | A | G | A | A | C | C | G | T | G | T | G | A | T | T | C | T | A | C | A | C | T | G | C | A | C |
| Sequenced event 13 | G | G | G | C | C | T | T | G | G | A | A | T | T | A | G | A | C | C | G | T | G | T | G | A | T | T | C | T | A | C | C | T | G | A | A | T | C | A |
| Sequenced event 14 | G | G | G | C | C | T | T | G | G | A | A | T | T | A | G | A | A | C | C | G | T | G | T | G | A | T | T | C | T | A | G | A | T | A | C | A | C | T |
| Sequenced event 15 | G | G | G | C | C | T | T | G | G | A | A | T | T | A | G | A | A | C | C | G | T | G | T | G | A | T | T | C | G | T | C | C | T | G | C | T | G | C |
| Sequenced event 16 | G | G | G | C | C | T | T | G | G | A | A | T | T | G | A | T | T | C | C | A | A | C | C | T | A | G | A | T | G | C | T | G | C | T | G | C | A | C |
| Sequenced event 17 | G | G | G | C | C | T | T | G | G | A | A | T | T | A | G | A | A | C | C | G | T | G | T | G | A | T | T | C | T | A | A | T | T | G | T | T | T | C |
| Sequenced event 18 | G | G | G | C | C | T | T | G | G | A | A | T | T | A | G | A | A | C | C | G | T | G | T | G | A | T | T | C | T | G | C | G | C | C | - | - | - | - |
| Sequenced event 19 | G | G | G | C | C | T | T | G | G | A | A | T | T | A | G | A | A | C | C | G | T | G | T | G | A | T | T | C | T | G | C | C | - | - | - | - | - | - |
| Sequenced event 20 | G | G | G | C | C | T | T | G | G | A | A | T | T | A | G | A | A | C | C | G | T | G | T | G | A | T | T | G | C | C | T | A | G | A | T | A | C | C |
| Sequenced event 21 | G | G | G | C | C | T | T | G | G | A | A | T | T | A | G | A | A | C | C | G | T | G | T | G | A | T | T | C | G | T | T | C | C | T | G | C | T | G |
| Sequenced event 22 | G | G | G | C | C | T | T | G | G | A | A | T | T | A | G | A | A | C | C | G | T | G | T | G | A | T | T | C | T | A | C | G | C | C | C | T | G | A |
| Sequenced event 23 | G | G | G | C | C | T | T | G | G | A | A | T | T | A | A | G | A | C | A | A | G | T | T | C | T | G | T | A | G | A | C | A | C | C | A | C | C | C |
| Sequenced event 24 | G | G | G | C | C | T | T | G | G | A | A | T | T | A | G | A | A | C | C | G | T | G | T | G | A | T | T | C | A | C | T | A | G | A | T | A | C | A |
| Sequenced event 25 | G | G | G | C | C | T | T | G | G | A | A | T | T | A | G | A | A | C | C | G | T | G | T | G | A | T | T | C | C | A | C | C | T | A | A | T | T | C |
| Sequenced event 26 | G | G | G | C | C | T | T | G | G | A | A | T | T | A | G | G | A | C | C | G | T | G | T | G | A | T | T | A | C | A | T | G | A | T | G | T | C | A |
| Sequenced event 27 | G | G | G | C | C | T | T | G | G | A | A | T | T | A | G | A | A | C | C | C | G | T | G | T | G | A | T | T | C | G | T | G | C | C | A | C | C | - |
| Sequenced event 28 | G | G | G | C | C | T | T | G | G | A | A | T | T | A | G | A | A | C | C | G | T | G | T | G | A | T | T | C | T | C | T | A | A | T | T | C | C | A |
| Sequenced event 29 | G | G | G | C | C | T | T | G | G | A | A | T | T | A | G | A | A | C | C | G | T | G | T | G | A | G | T | T | G | G | G | T | A | T | T | T | T | T |
| Sequenced event 30 | G | G | G | C | C | C | T | T | G | G | A | A | T | T | A | G | A | A | C | C | G | T | G | T | G | A | T | T | C | T | A | C | G | C | C | T | G | A |

Supplementary table 3: Protein and small molecule barcoded probes. All probes were purchased from IDT with high purity and desalting.

| Barcode name | Barcode sequence | Aptamer sequence | Aptamer reference |
|---|---|---|---|
| Adapter_barcode_Thrombin | GGGTTAAGCTATTGCTAACTGTAGTCCTAGTCTAGCTA | AGTCCGTGGTAGGGCAGGTTGGGGTGACT | Müller et al. (2008) J. Thromb. Haemost. |
| Adapter_barcode_BNP | GGGCGTAAACGTGAATGGGAGCCAATCATATACGTTGG | ATAACGACATCCGCCGGCACGAAGGGATCAAGTCGATAGG | Wang et al. (2014) Microchimica. Acta. |
| Adapter_barcode_cTnT | GGGCTTAAGTTCGGTCATTAACAGTGTCAATCTTGCCA | ATACGGGAGCCAACACCAGGACTAACATTATAAGAATTGCGAATAATCATTGGAGAGCAGGTGTGACGGAT | Sharma and Jang (2019) Sci. Rep. |
| Adapter_barcode_cTnI | GGGATTCAGATCTTTGACCGATCGTACTGACGTGTACG | AGTCTCCGCTGTCCTCCCGATGCACTTGACGTATGTCTCACTTTCTTTTCATTGACATGGGATGACGCCGTGACTG | Negahdary et al. (2017) Sens. Actuators B Chem. |
| Adapter_barcode_Serotonin | GGGACTAGATAAAAGGAAGGGAGCACAGTAACGTCGTT | CGACTGGTAGGCAGATAGGGGAAGCTGATTCGATGCGTGGGTCG | Nakatsuka et al. (2021) Anal. Chem. |

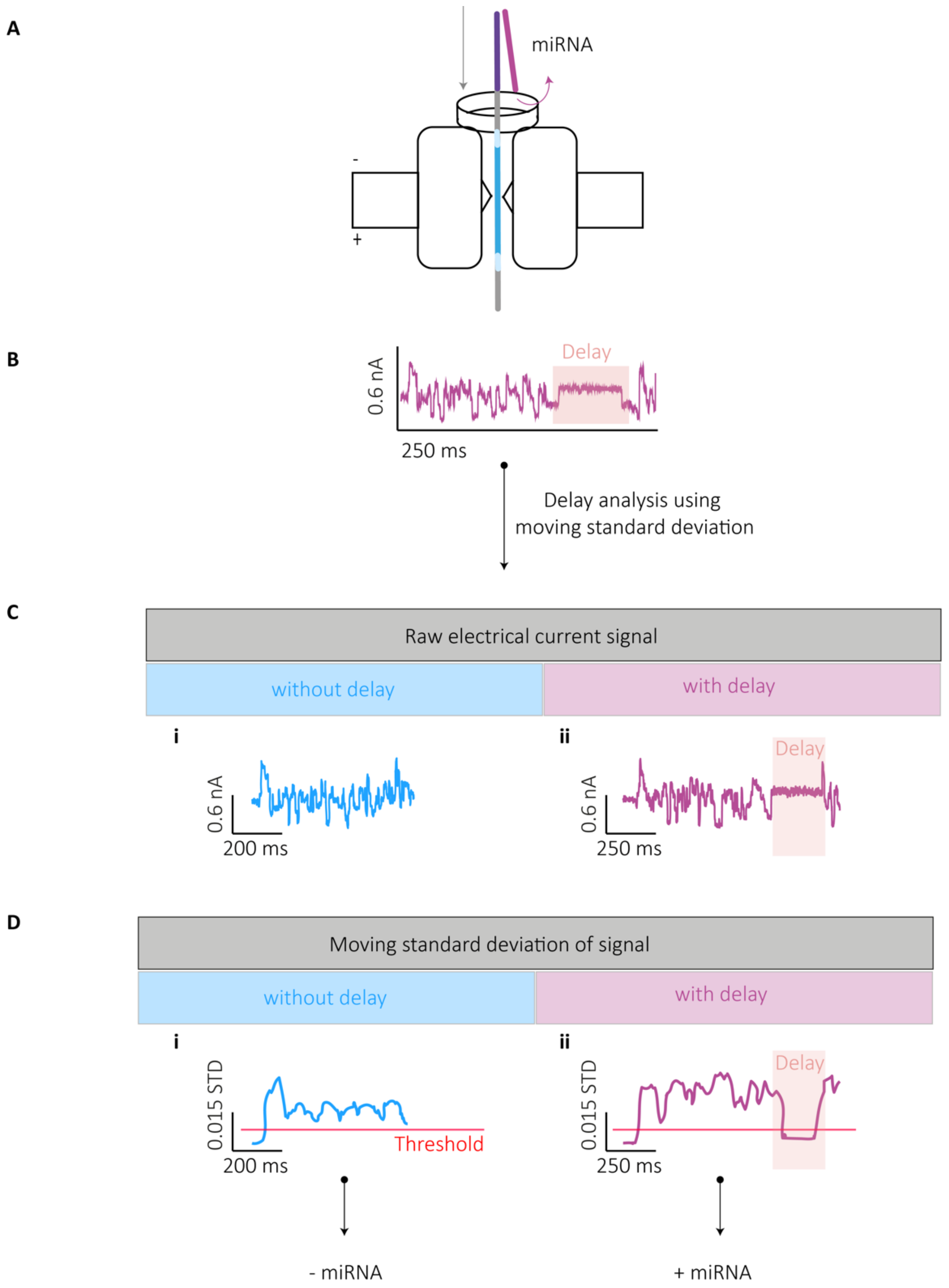


Supplementary figure 1: Delay detection method using moving standard deviation. A Illustration of barcoded probe translocation through a single nanopore. miRNA is bound to a section of the barcoded probe containing the complementary miRNA sequence. Double-stranded DNA / RNA is too large to translocate through the nanopore. The motor protein unzips the double-strand to allow the single-stranded barcoded probe to complete its translocation. B The period of strand unzipping can be observed in the electrical current as a "delay". C Example electrical current signals of barcode translocations without (i) and with (ii) delay periods. D Moving standard deviation of signal traces in part C. When the moving standard deviation drops below a threshold of 0.003 for a period of ≥10 bins (out of 75 bins total), the event is classified as delayed. Example values for events with (i) and without (ii) delay are shown.

A

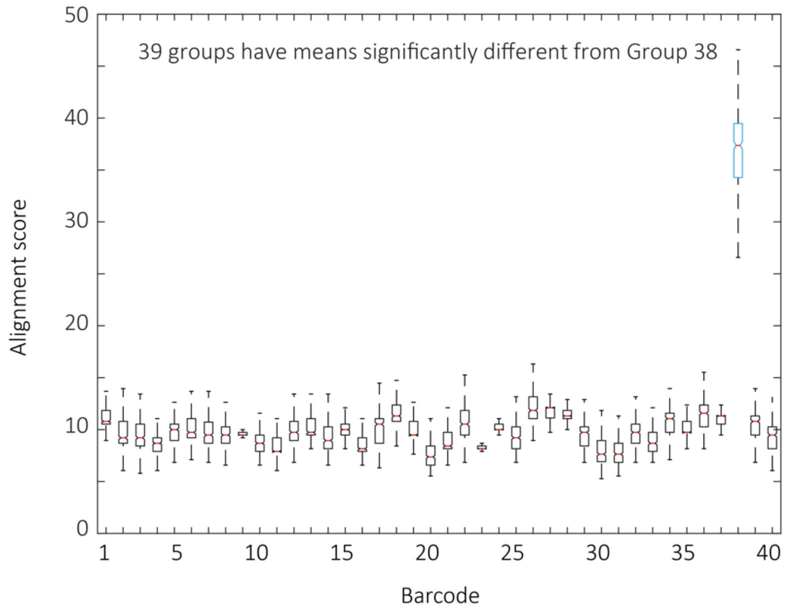


B

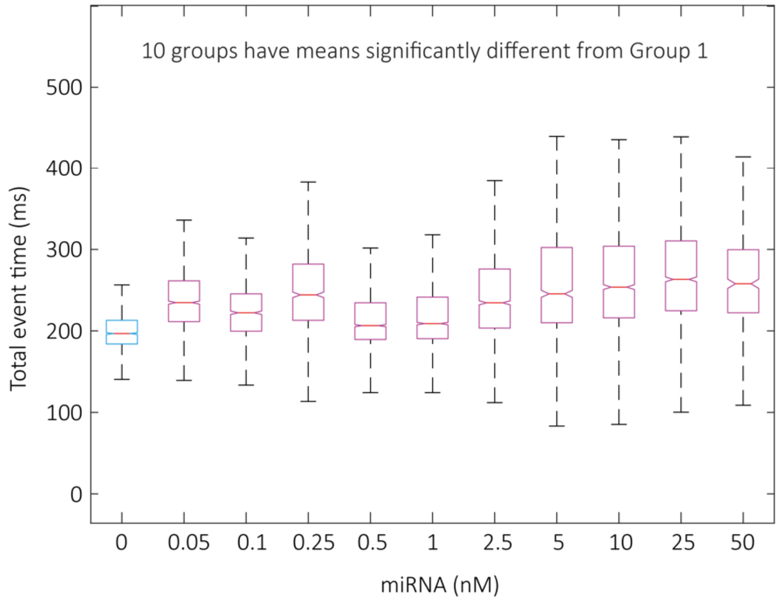


Supplementary figure 2: Statistical analysis of barcoded probe 38 events. A Alignment score of barcoded probe 38 events when aligned against all 40 barcoded probe sequences. Alignment score for the barcoded probe 38 sequence was significant from all other barcode sequences (ANOVA, $\alpha = 0.0001$, n=3). B Total event time (ms) of barcoded probe 38 events in the presence of various concentrations (0.05 to 50 nM) of its corresponding miRNA. Total event time was significantly increased in all conditions where miRNA was present when compared to control (ANOVA, $\alpha = 0.0001$, N=3, n=168-1845).

**A**

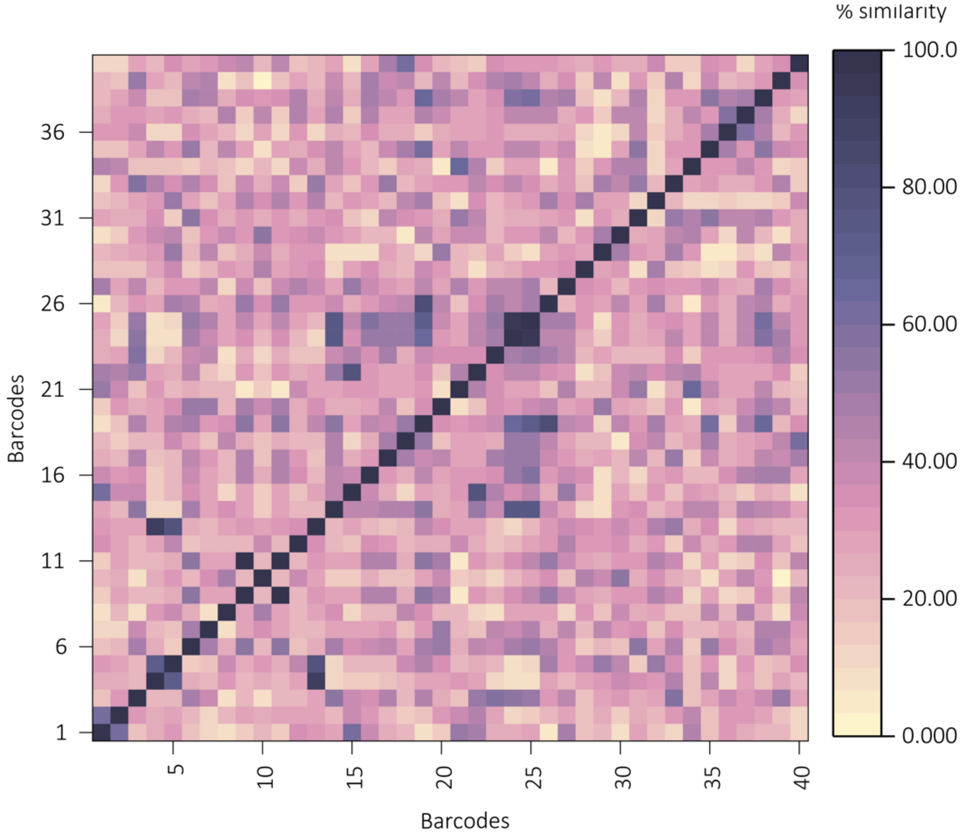


**B**

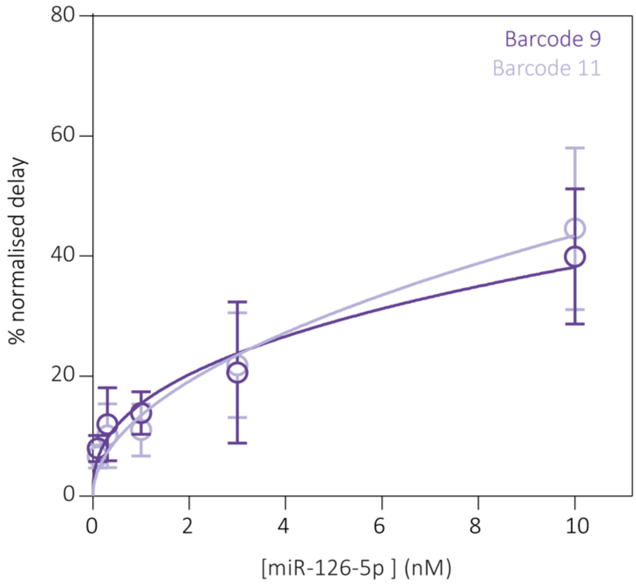


Supplementary figure 3: Sequence similarity of miRNAs in multiplexed experiments. A Confusion matrix of sequence similarity of miRNAs designed to bind to barcoded probes. Two barcoded probes (barcode 9 and barcode 11) were designed to bind the same miRNA (miRNA-126-5p). Two barcoded probes (barcode 24 and barcode 25) were designed to bind miRNAs with sequence similarity ≥95 % (miRNA-18a-5p and miRNA-18b-5p). Two barcoded probes (barcoded probe 4 and barcoded probe 13) were designed to bind miRNAs with sequence similarity 95 % > X ≥ 90 % (miRNA-30d-5p and miRNA-30e-5p). B Concentration- percentage delay event relationship of barcodes 9 and 11 (single barcode experiments). Both barcoded probes are designed to bind miRNA-126-5p. Barcode sequence had no significant effect on the % delay (N=3, n=3117-9310), meaning that both probes behave the same independent of the barcode sequence.

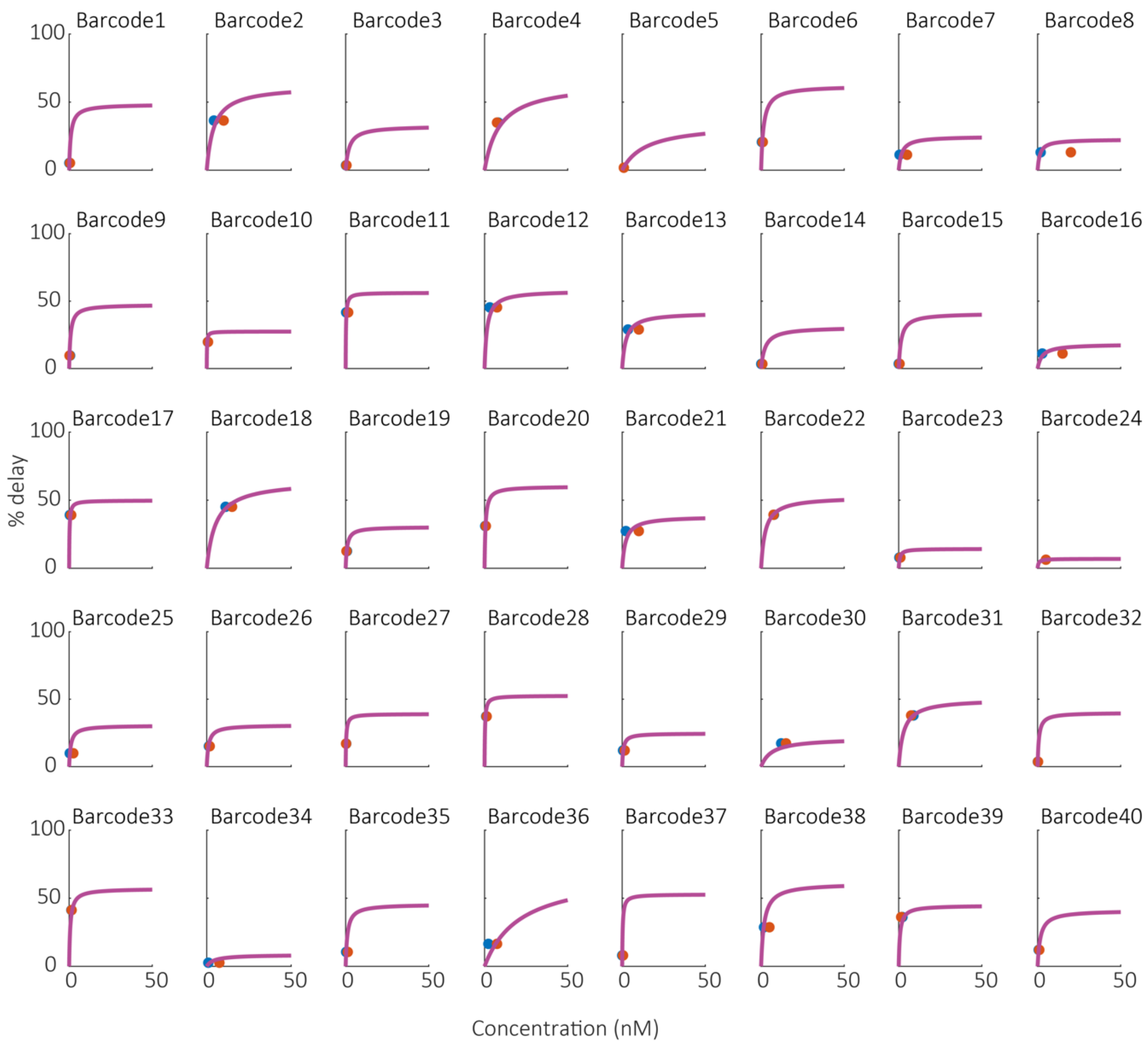


Supplementary figure 4: Individual barcoded probe concentration- delay relationships. Individual miRNA / barcoded probe titration curves are plotted with predicted (blue) and actual (red) values as determined in figure 4, (N=4, n=6189-71031).

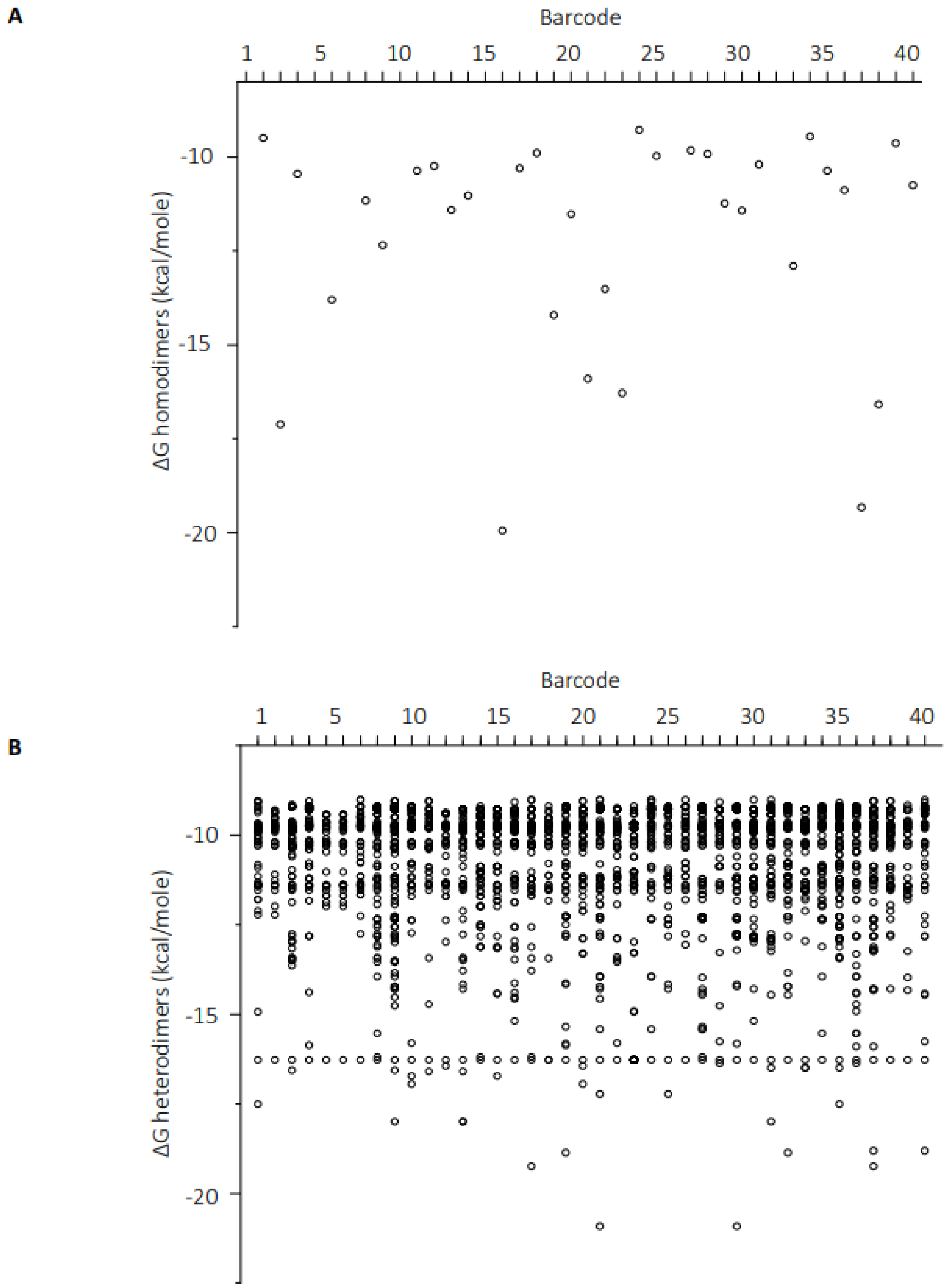


Supplementary figure 5: Prediction of barcoded probe interaction. Theoretical barcoded probe interactions were determined using an online web-app (Integrated DNA Technologies OligoAnalyser$^{TM}$ tool). Free energy of predicted homo-dimerisation of barcoded probes. Homo-dimers with ΔG ≤-9 kcal/ mol are shown. B Free energy of predicted hetero-dimerisation of barcoded probes. Hetero-dimers with ΔG ≤-9 kcal/mol are shown.

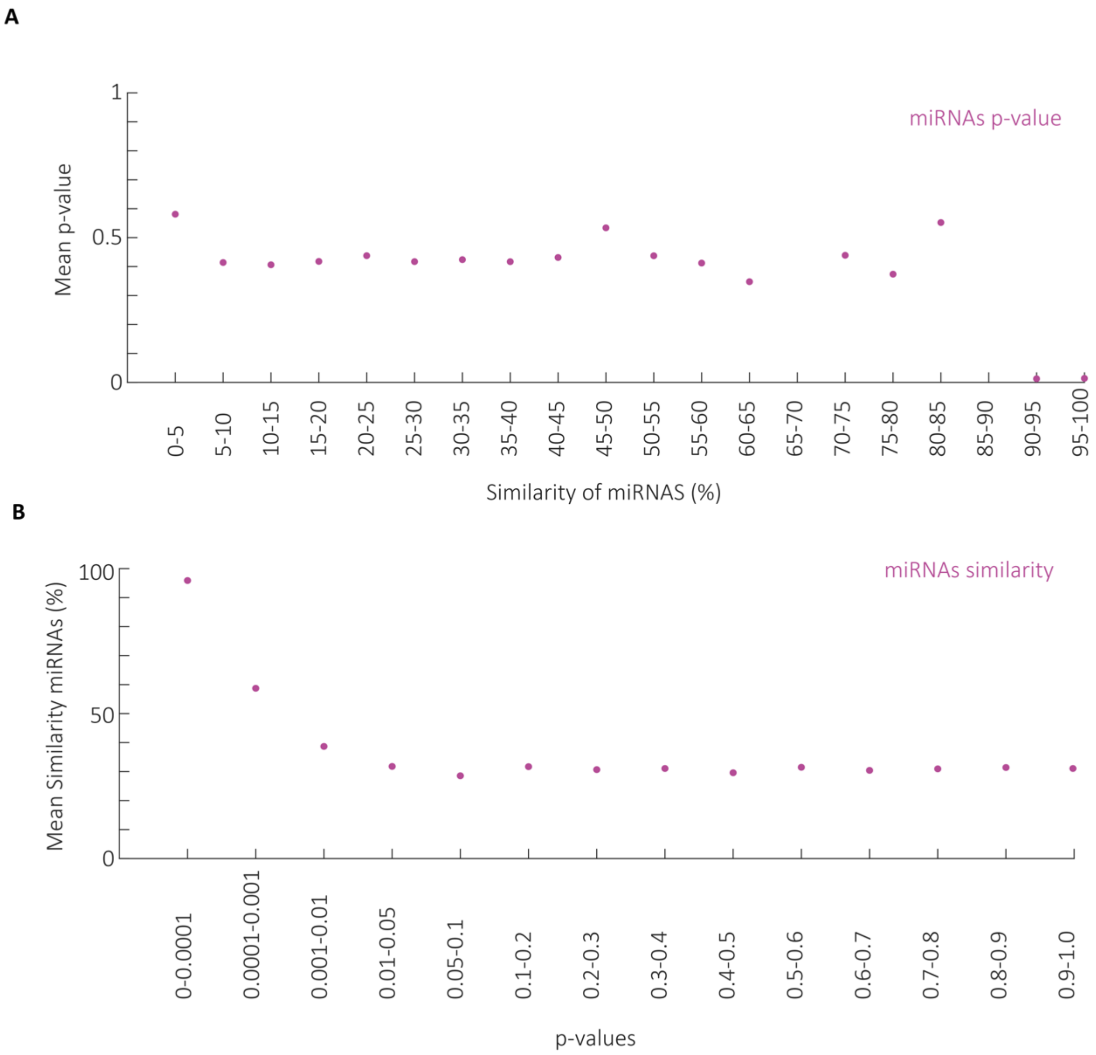


Supplementary figure 6: Probe selectivity of miRNAs. A Mean P-value (Students t -test), determined from change in percentage delay when single miRNAs were incubated with 40x barcoded probes, plotted against sequence similarity of miRNAs in each experimental condition (n=40). B Mean similarity score of miRNA sequences is inversely related to p-value (determined as above).

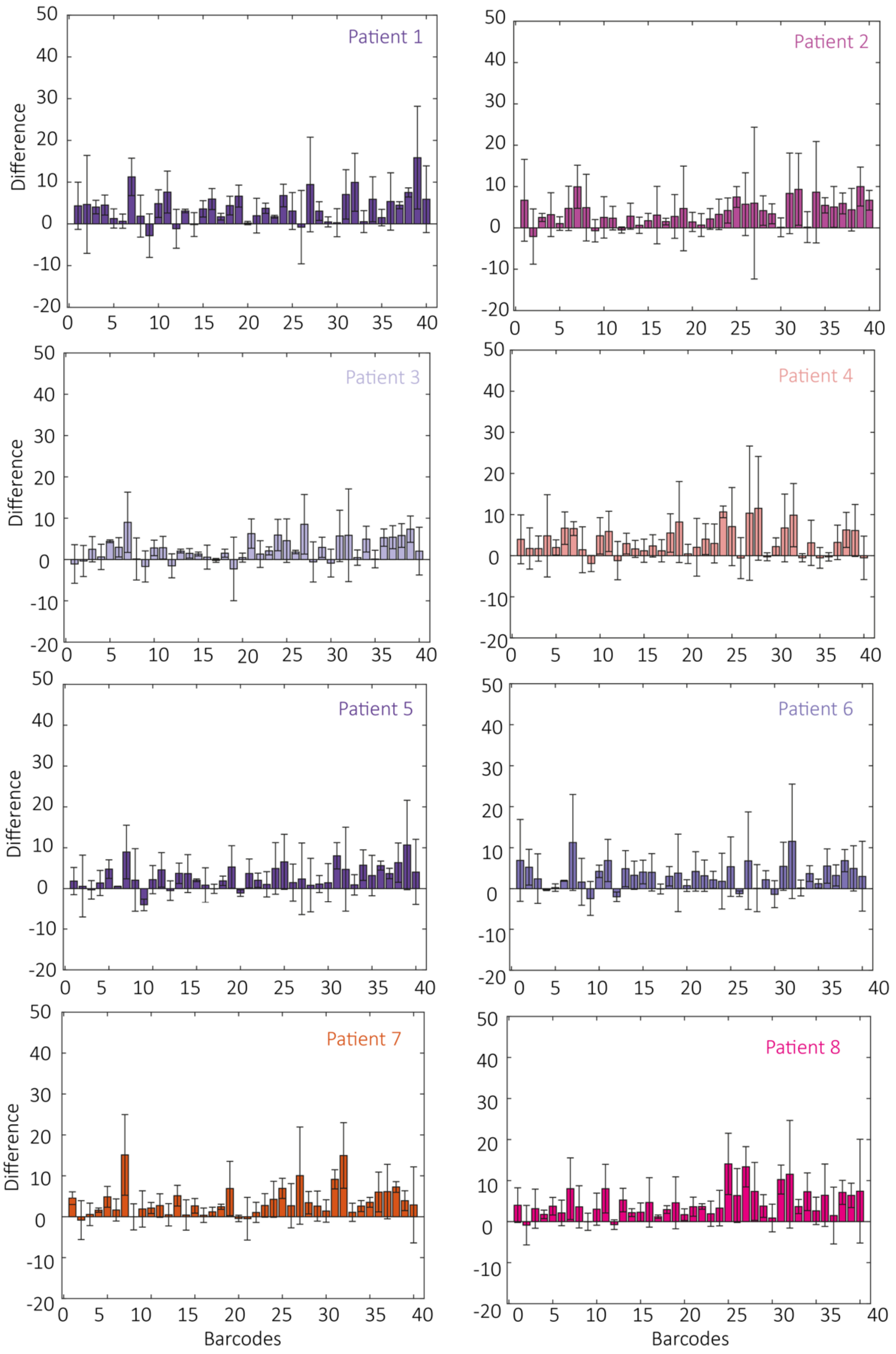


Supplementary figure 7: Analysis of patient samples. Change in percentage delay of barcoded probes due to addition of human serum, separated by participant is shown (N=3, n=17930-25776).

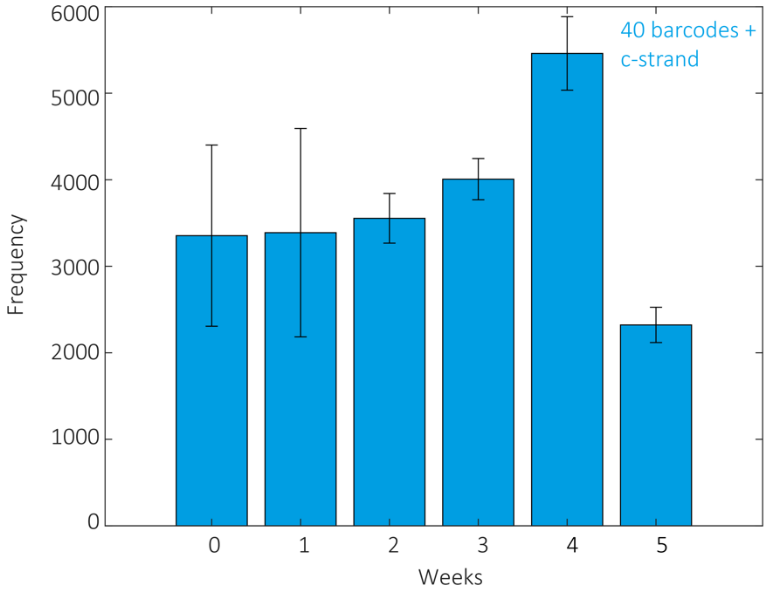


Supplementary figure 8: Lifetime analysis of frozen barcoded probes. A mix of 40 barcoded probes was stored at -20 $^{O}$C and assessed on a weekly basis for event frequency (N=2, n=2177-5760). Event frequency was reduced after the barcoded probe mix was stored for ≥5 weeks.

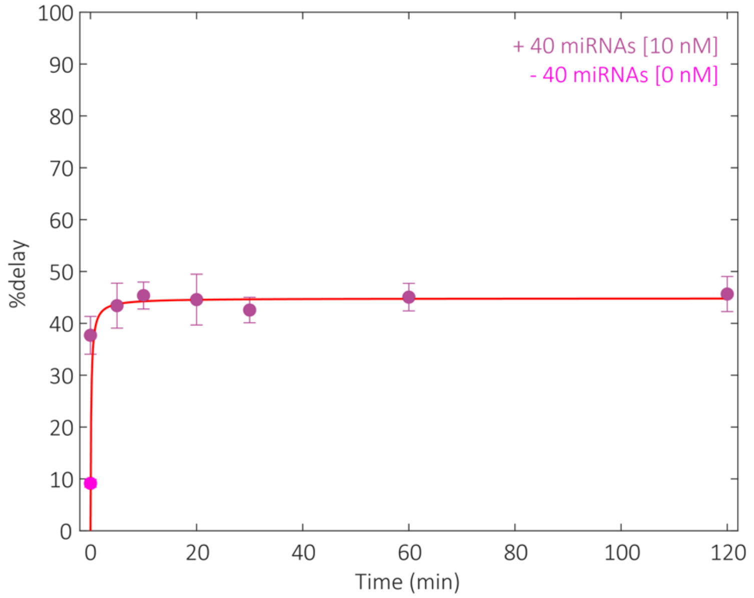


Supplementary figure 9: Barcoded probe binding to miRNAs equilibrates rapidly. A mixture of 40 barcoded probes and 40 corresponding miRNAs (10nM) were incubated for various time periods (0 – 120 minutes) and compared to the control (0nM miRNA). The mean percentage delay plateaued after 10 minutes (N=3, n=30902-43044).
